# Benchmarking cerebellar organoids to model autism spectrum disorder and human brain evolution

**DOI:** 10.1101/2025.05.14.653684

**Authors:** Davide Aprile, Oliviero Leonardi, Alessia Petrella, Davide Castaldi, Lorenza Culotta, Cristina Cheroni, Alessandro Valente, Matteo Bonfanti, Alessandro Vitriolo, Juan Moriano, Filippo Mirabella, Amaia Tintori, Cedric Boeckx, Giuseppe Testa

## Abstract

While cortical organoids have been used to model different facets of neurodevelopmental conditions and human brain evolution, cerebellar organoids have not yet featured so prominently in the same context, despite increasing evidence of this brain regions importance for cognition and behavior. Here, we provide a longitudinal characterization of cerebellar organoids benchmarked against human fetal data and identify at very early stages of development a significant number of dynamically expressed genes relevant for neurodevelopmental conditions such as autism and attention deficit hyperactivity disorders. Then, we model an ASD mutation impacting CHD8, showing both granule cell and oligodendrocyte lineages prominently affected, resulting in altered network activity in more mature organoids. Lastly, using CRISPR/Cas9 editing, we also model an evolution-relevant mutation in a regulatory region of the CADPS2 gene. We investigate the effect of the derived allele exclusive to Homo sapiens, identifying a rerouting of the CADPS2-expression in rhombic lip cells, coupled with a different sensitivity to hypoxia which in turn lead to a differential timing of granule cell differentiation.

**HIGHLIGHTS:**

- Longitudinal characterization of cerebellar organoids uncovers disorder related genes especially at early stages of development
- Mutation in CHD8 alter rhombic lip and oligodendrocytes differentiation via WNT pathway
- Rerouting of CADPS2 expression, delaying differentiation and migration, in recent human evolution

**IN BRIEF:** Aprile and colleagues longitudinally profiled cerebellar organoids, benchmarking them against a fetal human atlas and identified a highly dynamic expression of genes related to cognitive and behavioral disorders especially at early stages of differentiation. Organoids were used to model the impact of a high-penetrance mutation associated with autism spectrum disorder and a high-frequency derived allele in Homo sapiens predicted to have played a role in recent brain evolution.

## INTRODUCTION

Human cerebellar development is a particularly extended process that reaches completion around the first two years of postnatal life^1^. The complex cytoarchitecture of this brain region originates from two main proliferative pools of cells: the ventricular zone (VZ) and the rhombic lip (RL)^2^. While cell progenitors in the VZ gives rise to inhibitory neuron populations, including Purkinje cells, interneurons of the molecular layer, and deep inhibitory cerebellar nuclei, as well as most of the glial cells, the RL generates an entire array of excitatory neurons differentiating into granule cells (GCs), unipolar brush cells (UBCs), and deep excitatory cerebellar nuclei^3^. Altered development of these cell types hampers cerebellar function and morphology and causes major neurodevelopmental disorders as exemplified by spinocerebellar ataxia, medulloblastoma and other cerebellar malformations^4,5,6^.

More recent evidence has been pointing to an impaired cerebellar development as one of the best predictors of behavioral and cognitive disorders such as autism spectrum disorder (ASD), attention deficit and hyperactivity disorder (ADHD), and related psychiatric conditions^7,8,9,10,11,12,13^.

This reappraisal of the role of the cerebellum in cognition and social behavior, which significantly extends its historical preeminence in motility control, is emerging in parallel also from the context of human evolution. While the cortex has featured prominently in the anthropological and paleogenomic study of our most recent evolution, the cerebellum has thus far not received a commensurate attention, despite the modifications inferred from the fossil record point to its major contribution to the brain growth trajectory distinctive of our species^14,15,16,17,18^. Yet, despite the rising awareness that an increased focus on the cerebellum could greatly benefit progress in human neuroscience^19^, its modelling resource remains a largely undeveloped field. Animal models have contributed significantly to the general understanding of cerebellar development, but they have inherent limitations to probe human-specific spatio-temporal aspects of the cerebellar growth and maturation^2^, calling for a human-specific approach that build on the progress over the past decade with neural organoids, whose benchmarking ad standardization we extensively contributed to ^20,21^. To date, cerebellar organoid (CblO) generation is largely based - with few exceptions^22^ ^23^ - on the methodology set up by Muguruma and colleagues in 2015^24^ (eg. Nayler et al. 2021; Silva et al. 2021^25,26^). Most of these works exploit CblOs to study disorders such as medulloblastoma^27,28^, and spinocerebellar degenerative disorders^29^. Their potential to comprehensively model the cerebellum-specific developmental antecedents of behavioral and cognitive dysfunction is unexploited, as does their role in exploring recent human brain evolution at a higher resolution. In this work we begin to remedy this gap, hoping to shed light on some of the molecular and cellular foundations of the acquisition of higher-order cognitive functions in modern humans and their increased susceptibility to related neurodevelopmental disorders^30^. To do so, we benchmarked longitudinal transcriptomic profiles of CblOs from different control stem cell lines to reveal an early dynamic expression of genes related to ASD, ADHD and other psychiatric conditions. We exploited CblOs to model at this early stage of development the impact of a patient-specific and fully penetrant mutation in one of the most prominent ASD-causative genes, CHD8^31,32,33^, as well as the effect of a Homo sapiens-specific variant predicted to regulate CADPS2, a gene also associated with ASD and ADHD^34,35^ that lies in a region of positive selection and desert of introgression^16^.

## RESULTS

### Longitudinal characterization of cerebellar organoids (CblOs)

We differentiated CblOs from three control hiPSCs lines, CTL02A, CTL04E and CTL08A, adopting the experimental procedure illustrated in **Figure1A**, which builds on the seminal protocol published from Muguruma and colleagues in 2015^24^. The following protocol modifications were made: CblOs until 21 days of differentiation were kept at 3% oxygen (O_2_), then O_2_ tension was increased to 21% to mimic tissue vascularization occurring during cerebellar development^36^. Organoid’s growth tracked at early differentiation stages displayed a constant increase in dimensions at 3, 7 and 14 days, with no significant differences in cross-section area between the three lines measured as area under the curve among multiple differentiation rounds **(Figure 1B** and **Figure S1A-S1B**).

**FIGURE 1.**
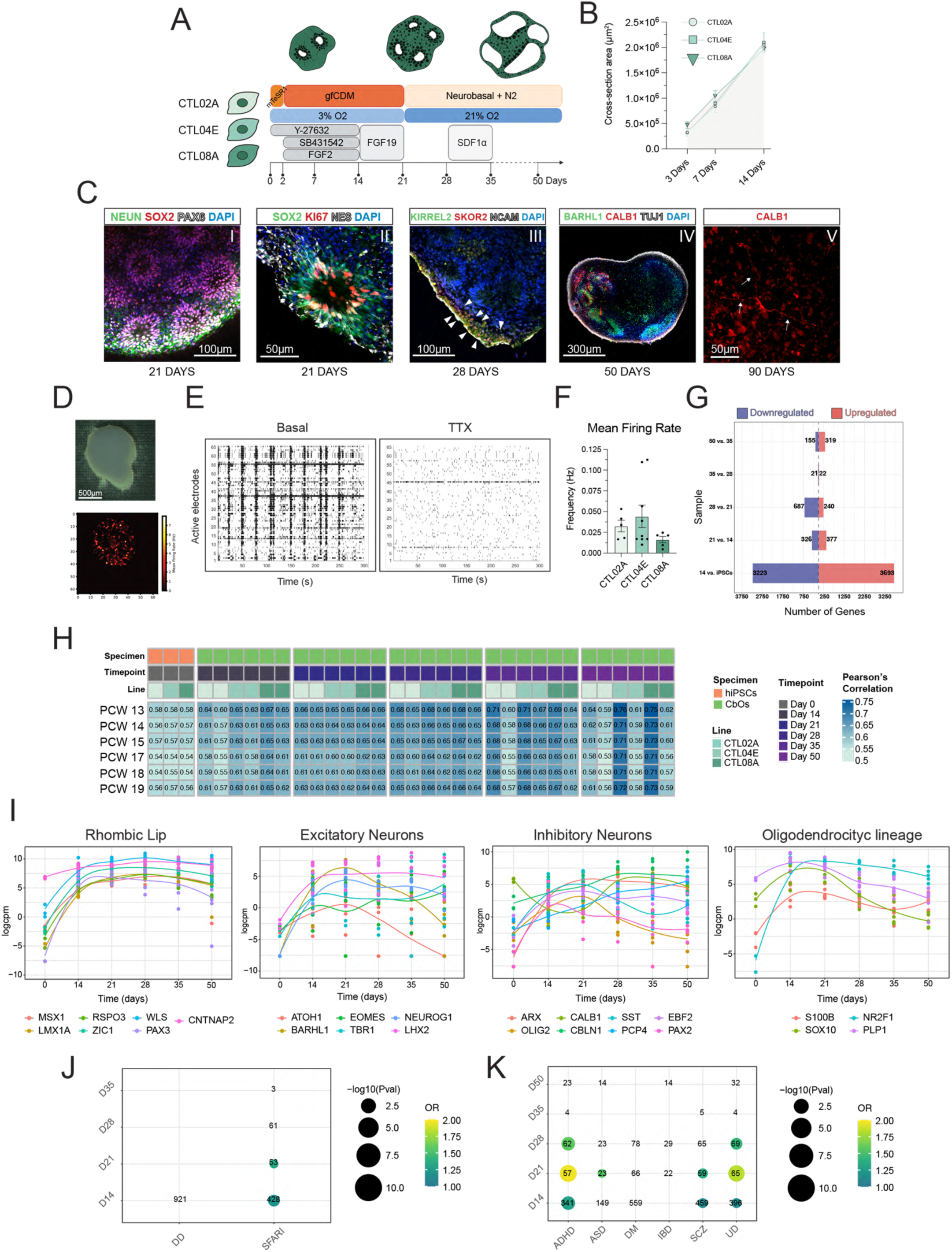
**Longitudinal characterization of cerebellar organoids (CblOs)** Schema showing the two male (CTL02A and CTL08A) and the female (CTL04E) hiPSC lines used in this study and the experimental design, culture media changes and small molecules/growth factors used in this work to differentiate CblOs at the different time points. (B) Time course showing CblO growth curves at 3, 7 and 14 Days of differentiation from the three control lines (CTL02A, CTL04E and CTL08A). Differences in organoids’ growth rate was measured by calculating the area under the curve generated plotting their major cross-section area along the timepoints. Measurements derives from three different differentiation batches. One way ANOVA with Tukey test CTL02A=1.24*107±9.49*105; CTL04E=1.29*107±1.81*106; CTL08A=1.35*107±6.38*105; CTL02A vs. CTL04E p>0.9999, CTL02A vs. CTL08A p=0.8902, CTL04e vs CTL08A p>0.9999. (C) Spinning-disk confocal images of immunocytochemistry followed by tissue clearing of CblOs at various differentiation stages. (I) neuroepithelial markers NES, SOX2 and PAX6 at 21 days; (II) co-labelling for SOX2 with the actively cycling cells marker KI67 and NES at 21 days; (III) neuronal adhesion molecule NCAM co-staining with CP and VZ markers (KIRREL2 and SKOR2 respectively), at 28 days. White arrowheads show SKOR2-positive cells at the organoid’s edge. (IV) RL and GCPs marker BARHL1 with early inhibitory neurons stained for CALB1 at d50. (V) Calbindin-positive neuron showed at 90 days of differentiation. Arrows indicates the arborization of the neuronal tree. Nuclei were stained with DAPI. (D) Representative image of a CblO from CTL04E at 130 days of differentiation on a MEA chip (top) and the active electrodes recorded from the very same organoid (bottom). (E) Raster plot showing basal network activity from and its suppression upon TTX administration. (F) Histograms showing burst frequency in CTL02A, CTL04E and CTL08A. Each dot represents the mean of the active electrodes for each organoid. Measures coming from two biological replicates, 4-8 differentiation batches CTL02A n=5, CTL04E n=9 and CTL08A n=5. One way ANOVA with Kruskal-Wallis test CTL02A vs. CTL04E p>0.9999; CTL02A vs. CTL08A p=0,7832; CTL04E vs CTL08A p=0,7197. (G) Butterfly plot comparing DEGs (FDR < 0.05, logFC > log2(1.5) as absolute value) from stage-wise differential expression analysis for each differentiation time point compared to the previous one from hiPSCs to CblOS at 50 days of differentiation in cerebellar organoid RNASeq dataset. Bar plots show on the x-axis the number of upregulated and downregulated genes for each comparison (as indicated on the y-axis). (H) Longitudinal correlation analysis of CblOs from 0 to 50 days of differentiation versus fetal cerebellar transcriptome at postconceptional weeks (PCWs) 13 to 15 and 17 to 19. The heatmap shows Pearson coefficient correlation values across stages. (I) Trend plots showing the expression levels (in logcpm) of cerebellar-specific genes in our bulk RNASeq dataset. Each plot reports key markers for the indicated lineages, and each dot represents a replicate derived from the three control lines (J & K) Overlap between the differentially expressed genes from stage-wise DEA (FDR < 0.05, logFC > 1 as absolute value) and gene-disorder knowledge bases. Panel J shows the overlap with SFARI genes and Development Disorder Genotype - Phenotype Database (DD); panel K represents the overlap with risk genes retrieved from GWAS catalogue for the following disorders: Attention Deficit Hyperactive Disorder (ADHD), Autism Spectrum Disorder (ASD), Diabetes Mellitus (DM), Inflammatory Bowel Disease (IBD), Schizophrenia (SCZ), Unipolar Depression (UD). Numbers represent shared genes and are shown for overlaps with odds ratio (OR) > 1, while dots are reported for overlaps having also a p-value < 0.05. Dot color represents OR values and dot size the p-value, as shown in the legend.

We characterized CblOs for the expression of cerebellar development-associated markers via immunocytochemistry followed by tissue clearing. Expression of neuroepithelial markers (SOX2 and PAX6) around ventricle-like structures reminiscent of the neural tube formation was identified at 21 days of differentiation (**Figure 1C**, **panel I)**. KI67-positive actively proliferating cells were found to reside within ventricle-like structures at the same differentiation stage, as showed in **panel II**. Later differentiation stages (28 days of development) revealed the presence of cerebellar markers characteristic of Cortical Plate (CP) and Ventricular Zone (VZ) development, such as KIRREL2 and SKOR2 (**panel III)**, and CALB1-positive early INs segregating from RL-like derivatives expressing BARHL1, as shown from staining at 50 days of differentiation **(panel IV)**. CALB1 was also used at 90 days of differentiation to track the morphological changes occurring in INs associated to cell maturation **(panel V)**. The increased arborization of INs was further confirmed by sparsely infecting hiPSCs with lentiviral particles carrying the expression of GFP upon the activation of the DLX1/2 promoter sequence and visualizing fluorescent cells at the same timepoint (**Figure S1C)**. Additionally, high-density MEA recordings were performed to reveal the functional properties of the neuronal network established within CblOs were performed **(Figure 1D)**. At 130 days, CblOs from the three control lines showed robust electrophysiological activity susceptible to the application of tetrodotoxin (TTX), a voltage-gated sodium channel blocker, indicating the occurrence of spontaneous firing events within the organoids **(Figure 1E and Figure S2D-S2E)**. Those events didn’t show any statistical difference across the three lines in terms of mean firing rate (MFR) and MFR extrabursts frequency **(Figure 1F, Figure S1F).**

To further characterize CblOs and identify the molecular changes occurring during their differentiation at salient timepoints associated with either the supplementation of specific small molecules, growth factors, culture media, and O_2_ tension, we performed a longitudinal transcriptional profiling by means of bulk RNA sequencing (bulk RNAseq). Starting from pluripotency (Day 0), we harvested CblOs at 14, 21, 28, 35 and 50 days of in culture and profiled each of the three lines in duplicate. To uncover the transcriptional changes along differentiation, we performed a time-wise differential expression analysis (DEA) by comparing each timepoint to the preceding one. This revealed a sizeable number of genes differentially expressed (DEGs) at early stages. This DEG set decreased (FDR < 0.05, FC > 1.5 as absolute value) as the differentiation progressed, indicating that most of the transcriptional modulation occurs at initial timepoints **(Figure 1G).** To better characterize these early expressed genes, we selected the most robustly up-regulated (FDR < 0.05, FC>2) until 28 days of differentiation, and characterized them via functional enrichment analysis. Gene Ontology (GO) terms retrieved from upregulated genes highlighted the occurrence of brain patterning events at early stages, followed by neuronal signaling and behavior-associated terms as differentiation proceeded (**Figure S1F**).

To benchmark CblOs against human cerebellar fetal tissue, we took advantage of a publicly available longitudinal profiling encompassing 6 different post-conceptional weeks (13-15,17-19^6^) and compared it to the gene expression dataset of our organoids. Our results showed an increased correlation in gene expression between our differentiating CblOs and the reference cerebellar transcriptome **(Figure 1H)**. At late stages, some organoids stochastically showed a reduced correlation value, due to an increased expression of choroid plexus-related markers such as transthyretin (TTR). At any rate, we established that this event, generated by an intrinsic ventral and caudalizing effect of the culture media, is not hampering the generation of cerebellar-specific lineages and the cell differentiation into mature neurons, as shown by immunostaining for BAHRL1 and TUJ1 (**Figure S1G**). We further examined if the expression of cerebellar markers driving tissue specification was temporally regulated along CblOs differentiation **(Figure 1I)**. We identified a robust modulation of genes associated with early Rhombic Lip (RL) development, such as LMX1A, RSPO3, PAX3, whose expression levels raised from around 14 to 21 days. This was followed by a decrease in the expression of late RL markers such as BARLH1 and ATOH1, and the peaking of more mature excitatory neurons (ENs)-associated transcription factors such as TBR1 and EOMES. Analogously, early VZ-derived inhibitory neurons (INs) markers such as PAX2 and OLIG2 peaked at 14 and 21 days of differentiation, respectively, followed by CBLN1, which reached its maximum expression at 28 days of differentiation along with PCP4, associated with Purkinje cells (PCs) differentiation, and whose expression trend constantly raised along CblO maturation. Lastly, we also observed a dynamic expression of genes associated with the oligodendrocyte lineage, such as SOX10 and S100B, both peaking at early differentiation stages. The expression of these genes was inversely modulated at late timepoints, with the decrease of SOX10 and the rise of S100B, likely indicating an exhaustion of the pool of oligodendrocyte precursors (OPCs) and the establishment of a more mature set of oligodendrocytes (OLs).

Finally, we assessed the relevance of changes in expression patterns along organoid differentiation to investigate their use in modeling the molecular bases of neurodevelopmental and psychiatric conditions. For this we performed an overlap analysis between the DEGs retrieved from the time-wise DEA (FDR < 0.05, log2FC > 1 as absolute value) and either genes associated with neurodevelopmental diseases of high-penetrance or genes included in the EBI GWAS catalog^37^ of variants associated with neurodevelopmental conditions presenting behavioral and cognitive dysfunction **(Figure 1J-1K).** We identified between 14 and 21 days of organoids’ differentiation, a significant overlap between DEGs from CblOs and genes listed within the SFARI database (https://gene.sfari.org). Similarly, genes associated to ADHD, unipolar disorder (UD), schizophrenia (SCZ) and ASD were found to be differentially expressed in CblOs between 14 and 20 days of differentiation.

### ASD-associated mutation in CHD8 induces CblOs transcriptional changes at early stages of differentiation

To demonstrate that CblOs provide a valuable tool to understand the molecular mechanisms at the basis of cerebellar dysfunctions, we resorted to CHD8 as a paradigmatic gene with well-established associations to neurodevelopmental disorders (NDDs). Specifically, we used a transgenic line obtained through CRISPR/Cas9 editing of human embryonic stem cells (hESCs) H9 (CHD8^+/+^), carrying in hemizygosity the single nucleotide variant (SNV) E1114X, described in Villa et al. 2021^38^. This isogenic CHD8^+/E1114X^ mutant, constitutively expresses the enhanced version of the green fluorescent protein (eGFP) **(Figure 2A)**. As our characterization of CblOs described above was performed on hiPSCs-derived organoids, we first compared CblOs differentiated upon either hiPSCs or CHD8^+/+^ (hESCs). CblOs derived from both CTL04E and CHD8^+/+^ lines showed similar growth rates as illustrated by transmitted light images and growth curves comparison (**Figure S2A -S2C**). Additionally, RL- and VZ-derived cell markers (BARHL1 and CALB1), similarly distribute in CblOs from both lines and MEA recordings comparing CblOs derived from hESCs and the three hiPSCs lines used in the previous section at 130 days didn’t show differences in MFR **(Figure S2D - S2E**). Lastly, comparison of the transcriptomic profile from CTL04E and CHD8^+/+^ bulk RNAseq at 14 and 21 days highlight strong correlation across both timepoints, indicating the interchangeable use of these cell lines at the developmental stage of interest for our analyisis.

**FIGURE 2.**
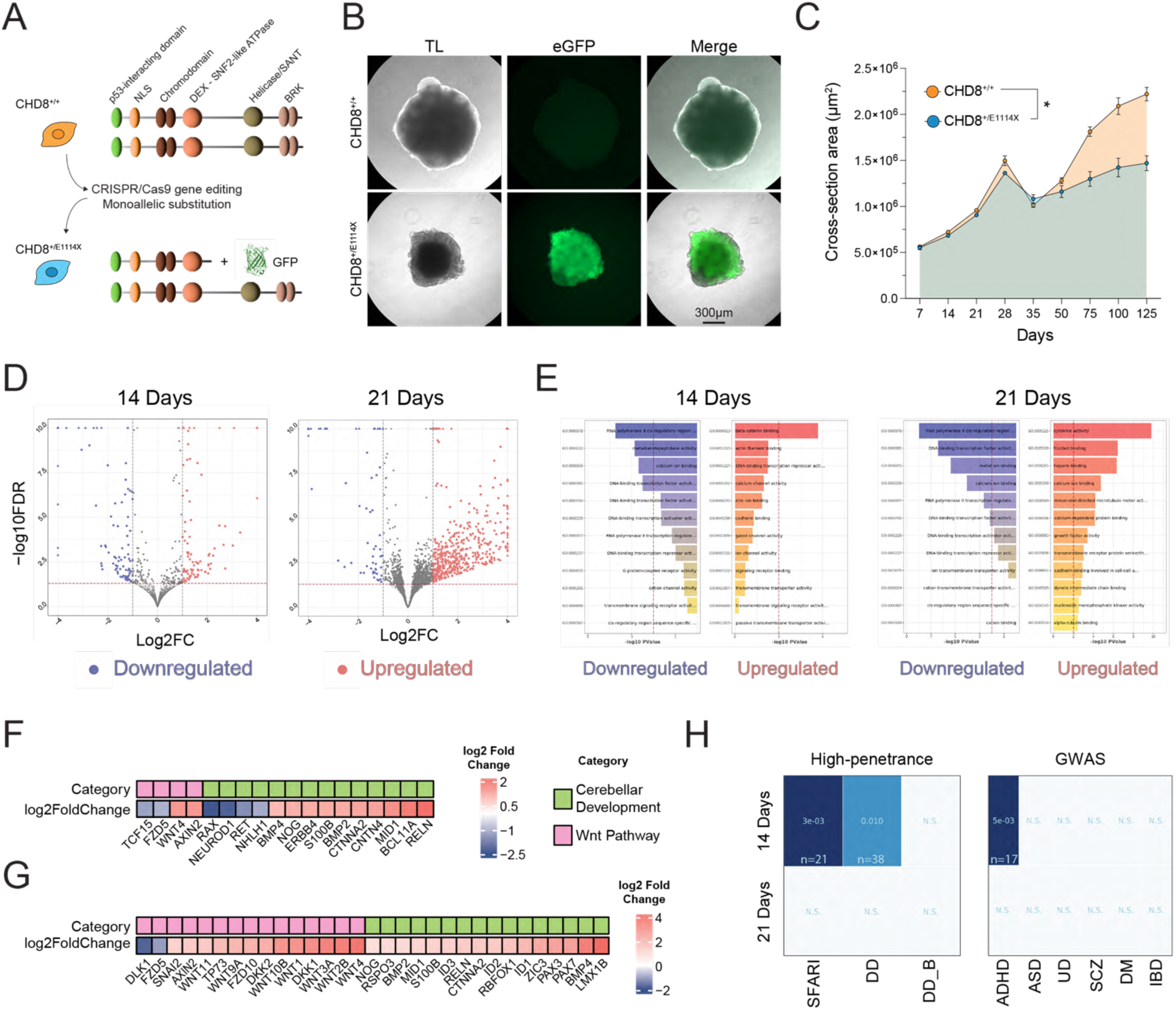
Mutations in CHD8 induce CblOs transcriptional changes at early stages of differentiation. **(A)** Schematic showing the H9 hESCs line (CHD8^+/+^) engineered through CRISPR/Cas9 editing to insert in heterozygosity the E1114X mutation along with eGFP expression. eGFP cartoon from https://www.rcsb.org/structure/1EMA **(B)** Representative transmitted light images from CHD8^+/+^ and CHD8^+/E1114X^ -derived CblOs at 50 days of differentiation. eGFP expression of CHD8^+/E1114X^ is showed in green. **(C)** Time course showing CblOs growth curves from 7 to 125 days of differentiation from CHD8^+/+^ and CHD8^+/E1114X^. Measurements derives from 4 to 10 organoids at least from four differentiation batches. **(D)** Volcano plots representing the results of the differential expression analysis for CHD8^+/E1114X^ *vs.* CHD8^+/+^ CblOs at 14 (left panel) and 21 (right panel) days of differentiation. 2-4 samples were analyzed per each genotype at the different timepoints. Results are reported for the pool of tested genes as – log10FDR (y-axis) and log2FC (x-axis). Genes identified as significantly modulated (FDR < 5% and absolute log2Fc >1) are shown in red for upregulated genes and in blue for downregulated ones. **(E)** Results of gene ontology enrichment analysis performed on the Molecular Function domain of GO on the DEGs, split in upregulated (FDR < 0.0.5, log2FC > 1) and downregulated (FDR < 0.05, log2FC < -1). Bar plots show TopGO p-value for the top 12 most significant categories; red dotted line represents a p-value threshold of 0.01. **(F & G)** Heatmap showing genes associated with cerebellar patterning (Cerebellar Development) and the WNT cascade (WNT Pathway), differentially expressed in CHD8^+/+^ vs. CHD8^+/E1114X^ - derived CblOs. **(H)** Overlap between the differentially expressed genes in CHD8^+/+^ vs. CHD8^+/E1114X^ from DEA between and gene-disorder knowledge bases. The left panel shows the overlap with SFARI (n = 21) and DD (n = 38) genes, whether the right panel represents the overlap with risk genes retrieved from GWAS catalogue for ADHD (n = 17), ASD, DM, IBD, SCZ, UD.

We then tested CHD8^+/E1114X^ cells as model recapitulating CHD8 haploinsufficiency *in vitro*. CHD8 expression was found to be reduced in CHD8^+/E1114X^ hESCs compared to controls in immunocytochemistry experiments quantifying CHD8 staining within the nuclear compartment **(Figure S3A-S3B)**. Likewise, quantification of CHD8 expression from western blot experiments performed on cytosolic/nuclear fractions **(Figure S3C- S3D)** showed a reduction of CHD8 levels in CHD8^+/E1114X^ compared to CHD8^+/+^ in the nucleus. CblOs were then differentiated from both CHD8^+/+^ and CHD8^+/E1114X^, with mutants exhibiting a difference in morphology during development, showing rough edges and a significantly smaller size in comparison to their relative controls **(Figure 2B)**. This phenotypic difference was corroborated by a growth curve analysis where we measured the cross-sectional area of CblOs in a time course spanning from 7 to 125 days of differentiation. The analysis revealed a significant reduction in the area under the curve (AUC) of CHD8^+/E1114X^ compared to CHD8^+/+^ **(Figure 2C, Figure S3E)**. This result is coherent with findings in conditional *Nestin-Cre/Chd8* haploinsufficient mice showing reduced cerebellar size ^39^ but it is in contrast with the overall macrocephalic phenotype observed in CHD8^+/E1114X^ patients and recapitulated in cerebral organoids^38^, pointing to a significant region-specific effect.

In light of the well-established role of CHD8 in the regulation of gene expression, we sought to investigate how the E1114X variant could impact on the transcriptomic landscape of developing CblOs. For this, we performed bulk RNAseq experiments at the two timepoints enriched in DEGs related to the etiology of cognitive and behavioral disturbances in patients, namely 14 and 21 days of differentiation, for both CHD8^+/+^ and CHD8^+/E1114X^ -derived CblOs. We compared for each time-point CHD8^+/E1114X^ to CHD8^+/+^ by differential expression analysis (**Figure 2D**). At 14 days, we found a mild modulation of the transcriptional landscape, with a similar effect on up-regulated and down-regulated genes (respectively 92 *vs.* 95 DEGS, FDR < 0.05 and logFC > 1 as absolute value). By contrast, at 21 days, the transcriptional effect of CHD8 mutation turned out to be much more pervasive, with a striking unbalance between the up- and the down-regulated genes (479 *vs.* 67 DEGs respectively). Functional enrichment analysis for the Molecular Function domain of Gene Ontology (GO) was then performed separately for down-regulated and up-regulated genes for both time-points. At 14 days, we found a significant downregulation of GO terms associated with RNA polymerase activity and DNA transcription, in line with the known role of CHD8 in regulating these processes^40,41^. In parallel, we identified an upregulation of genes mediating the activity of β-catenin, an established molecular CHD8 interactor^42,43^, having a role in the context of neuronal differentiation^44,45^ and more specifically during cerebellar development in mice^46,47^ (**Figure 2E, left)**. At 21 days, along with the downregulation of GO terms associated to transcriptional activity already present at earlier stages, we found an upregulation of genes associated with alpha tubulin binding, a feature of many chromatin modifier associated with ASD^48^, in addition to GO terms associated with growth factors activity and signal transduction processes, indicating a role for CHD8 in tissue patterning and cell differentiation during CblOs development **(Figure 2E, right).** The correlation between CHD8 activity and CblOs differentiation is further corroborated by the manually curated set of DEGs presented in **Figure 2F and Figure 2G**, where a disruption of genes orchestrating cerebellar development is evident, alongside a marked change in the expression of genes related with the WNT/β-catenin pathway. We then overlapped for both the timepoints the list of DEGs with the dataset of high-penetrance and GWAS knowledge-based genes associated with neurodevelopmental conditions presenting cognitive and behavioral disturbances used above (**Figure 1J-1K**). A significant overlap with high-penetrance genes was found at 14 days with the SFARI collection and the Development Disorder Genotype-Phenotype Database (DD) as well as with genes associated with ADHD through GWAS (**Figure 2H**). Some of these genes have an established role in cerebellar development, such as RELN^49,50^, GRID2^51^ and MID1^52^.

### scRNAseq analysis shows alterations in cell population composition in CHD8^+/E1114X^ -derived CblOs

To investigate cell type-specific effects of CHD8 depletion, we performed single-cell RNA sequencing (scRNAseq) of CHD8^+/+^ and CHD8^+/E1114X^ CblOs at 21 days of differentiation, encompassing around 18,000 high quality cells. After normalization and dimensionality reduction, the UMAP showed a clear segregation of the two conditions, mostly occupying different regions of the UMAP space **(Figure 3A)**, consistently across replicates (Figure S4A). Unbiased clustering followed by cell type annotation by known population markers genes led to the identification of 16 clusters **(Figure 3B)**. To quantify the different distribution of the two genotypes in the identified cell populations, we performed a differential abundance analysis by Milopy^52^ that confirmed a major impact of the CHD8 mutation CblOs composition (**Figure 3C**). Specifically, we identified within the dataset five neural stem cells (NSCs) clusters, expressing proliferation genes attributable to different stages of the cell cycle **(Figure S4B).** Among those five clusters, three were exclusively populated by the CHD8^+/+^ genotype, as with most of the postmitotic neurons, both excitatory (ENs) and inhibitory (INs) (Figure 3D). Consistent with their early stage of differentiation, the ENs annotation captured both granule cells progenitors (GCPs) markers (EOMES) and cerebellar excitatory interneurons markers (LHX2, TBR1) expressing cells, whereas INs expressed canonical markers like GAD2 and DLX1/2/5/6. In contrast to controls, where RL-associated cell types were clearly defined into precursors and differentiated cells, CHD8^+/E1114X^-derived organoids generated a continuous gradient composed of early (SOX2+) and late (PAX6+) RL-like cells, along with BARHL1+ GCPs and immature granule neurons (iGNs; expressing DCX and NEUROG1). Two additional clusters of proliferating ciliated cells identified as ependymal cells expressing dorsal (ZIC1) or ventral markers (ARX) were associated exclusively with CHD8^+/E1114X^. Last, despite an almost equal representation in terms of cell numbers within the oligodendrocyte precursors (OPCs) lineage, characterized by the expression of makers such as NES and PDGFRA/B, mature oligodendrocytes (OLs) expressing CNP, PLP1, SOX10 and S100B were found to be enriched exclusively in CHD8^+/E1114X^ **(Figure S4C)**. To investigate whether the modifications in cell type composition were the result of divergent developmental trajectories, we calculated diffusion maps separately for CHD8^+/+^ and CHD8^+/E1114X^ cells. This representation highlighted for CHD8^+/+^ cells a distinct transcriptomic difference between NSC and both ENs and INs neurons, clearly widespread across the two diffusion components (DC) 1 and 2 (**Figure 3E**). By contrast, mutant cells displayed an aligned clustering prevalently along DC2 (**Figure 3F**), corroborating the idea of an altered developmental trajectory in CHD8^+/E1114X^ CblOs. Looking more closely at the most divergent lineages, we experimentally confirmed that BARHL1 exhibits increased expression in mutants compared to controls, by immunostaining quantification (**Figure 3H-3I**) and qPCR (**Figure S4D**). Likewise, we confirmed that the number of cells expressing S100B was enriched in CHD8^+/E1114X^ compared to controls (**Figure 3J**), quantifying S100B expression *via* immunostaining (**Figure 3K-3L**) and qPCR (**Figure S4E**). We then investigated the possibility that β-catenin alterations, as suggested by bulk transcriptomics, could have influenced the acquisition of the altered cell fate revealed by scRNAseq for CHD8^+/E1114X^. Although quantification of total β-catenin levels through western blot on CblOs lysates did not reveal any changes at the protein level (**Figure 3M – 3N**), we computationally inferred transcription factor (TF) activity to probe β-catenin activity at downstream targets, and this resulted in the identification of a large group of cells of the CHD8^+/E1114X^ genotype with high levels of imputed β-catenin activity (**Figure 3O**). In particular, RL clusters and iGNs along with the oligodendrocyte lineage were the most affected. Quantitative PCR (qPCR) on CblOs at Day 21 **(Figure S4F**), showed, for CHD8^+/E1114X^ compared to controls, an increased expression of AXIN2, DKK1 and SP5, all transcriptional targets of β-catenin associated with the repressor activity of this molecular cascade^53–55^. We interpret this experimental outcome as the effect exerted by a constantly active β-catenin downstream mechanism failing to establish a functional negative feedback in CHD8^+/E1114X^ CblOs, coherently with previously published literature connecting an overactivation of the WNT pathway with a reduction in cerebellar size along with altered cell differentiation^56^.

**FIGURE 3.**
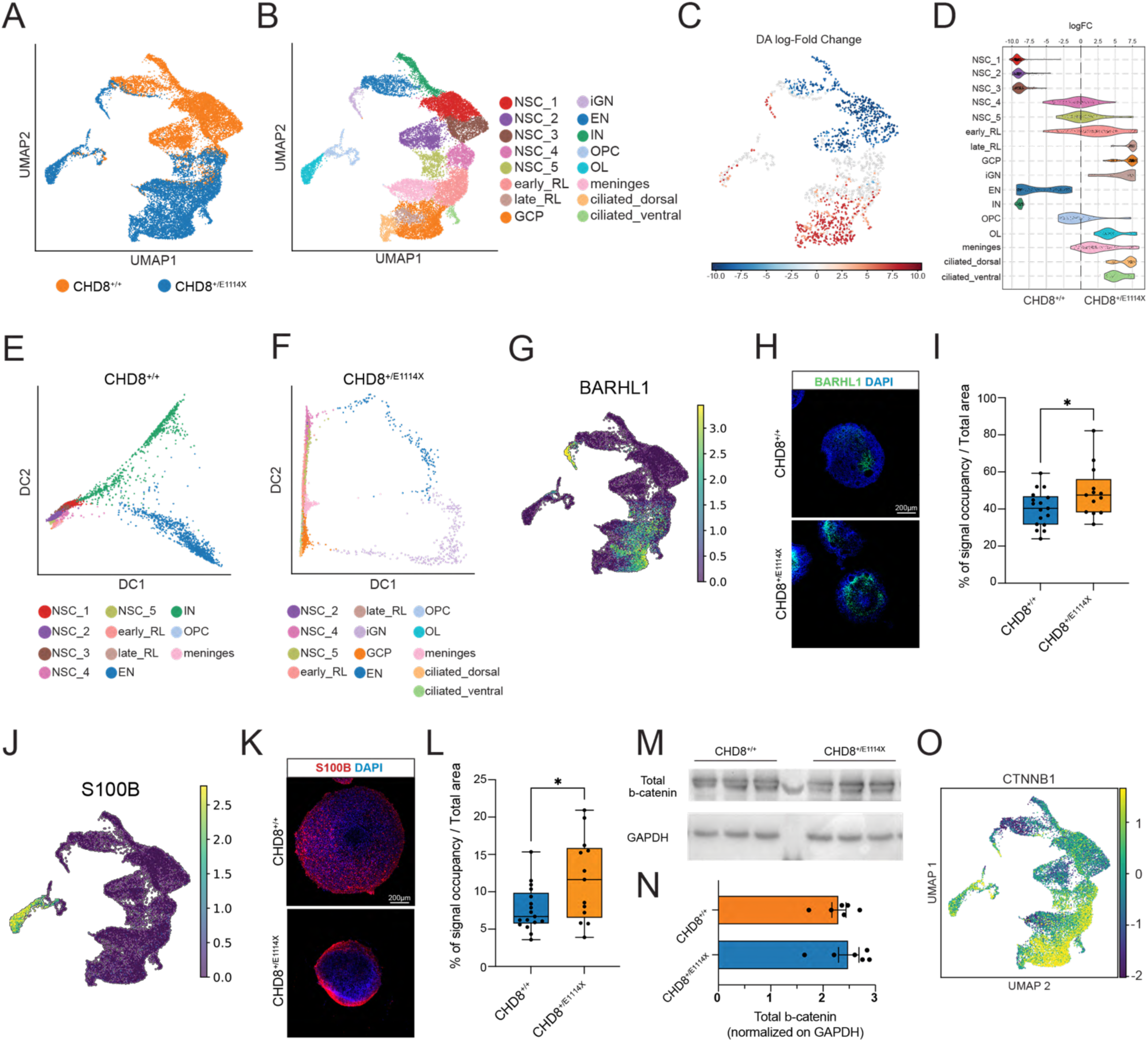
scRNAseq analysis shows alterations in cell population composition in CHD8^+/E1114X^ - derived CblOs. **(A-B)** Uniform manifold approximation and projection (UMAP) calculated on the scRNAseq data from CHD8^+/E1114X^ and CHD8^+/+^ CblOs at day 21. Each dot is colored according to genotype in (A) CHD8^+/+^ = 9806 cells (orange), CHD8^+/E1114X^ = 10400 cells (blue), and annotated cell type in (B). Annotation performed on Leiden clusters based on known cell markers retrieved the following cell populations: Neuronal Stem Cells (NSCs), early and late rhombic lip (early_RL, late_RL), granule cells progenitors (GCP), immature granule neurons (iGN), excitatory neurons (EN), inhibitory neurons (IN), oligodendrocyte precursors (OPC), oligodendrocytes (OL), meninges, ciliated dorsal and ventral. **(C)** Results of differential abundance analysis by Milopy comparing CHD8^+/E1114X^ to CHD8^+/+^ and visualized on the UMAP. Color-code indicates the enrichment in mutant (red) or wild-type (blue) and is shown only for significantly enriched neighborhoods (FDR < 0.01). (**D)** Violin plots showing for each neighborhood the fold-change calculated by Milopy; each neighborhood is assigned to the most abundant cell type. **(E & F)** 2D diffusion maps calculated separately for CHD8^+/+^ and CHD8^+/E1114X^ cells (left and right panel respectively) and colored according to the annotated cell type. **(G)** UMAP showing the expression levels of BARHL1 in the scRNAseq dataset. **(H**) Representative spinning-disk confocal microscope image showing 21 days CblOs stained for BARHL1 (green), and DAPI (blue). **(I)** Box plot showing the quantification of fluorescence signal in 21 days organoids labelled for BARHL1 and expressed as % of fluorescence signal occupancy over the organoids’ area measured over different z-stacks. Unpaired t test, p = 0,0261 (*). CHD8^+/+^ n=17, 40,19 ± 2,355; CHD8^+/E1114X^ n =13, 49,28± 3,793. Results from 4 independent experiments. **(J)** UMAP showing the expression levels of S100B in the scRNAseq dataset. **(K)** Representative spinning-disk confocal microscope image showing 21 days CblOs stained for S100B (red), and DAPI (blue). **(L)** Box plot showing the quantification of fluorescence signal in organoids at 21days labelled for S100B and expressed as % of fluorescence signal occupancy over the organoids’ area measured over different z-stacks. Unpaired t test, p = 0,0251 (*). CHD8^+/+^ n=17, 7,709± 0,7260; CHD8^+/E1114X^ n =13, 11,66±1,550. Results from 4 experimental replicates. **(M & N)** Western blots showing overlay for active and total b-catenin in CblOs at 21 days of differentiation from both CHD8^+/+^ and CHD8^+/E1114X^. In (N) histograms show total b-catenin as ratio normalized on GAPDH expression. Data from 3 independent organoids preparations. CHD8^+/+^ = 0,304±0,02302; CHD8^+/E1114X^ = 0,2446±0,01348. Mann-Whitney, p=0,0317 (*). **(O)** UMAP showing CTNNB1 activity as calculated by transcription factor analysis showing CTNNB1activation in CHD8^+/+^ and CHD8^+/E1114X^ within the dataset.

### CHD8^+/E1114X^ -derived CblOs show functional alterations of network activity at late developmental stages

The striking differences in cell type abundance between CHD8^+/E1114X^ and controls at 21 days of differentiation prompted us to investigate the functional impact of these cellular composition changes at advanced developmental stages, focusing on the two main lineages impacted by CHD8 mutation: RL-derived cells and oligodendrocytes. Our starting point was a functional enrichment analysis on genes differentially expressed between CHD8^+/+^ and CHD8^+/E1114X^ at 21 days of development: this analysis highlighted many terms associated with neuronal excitatory transmission and synaptic compartment maturation in CHD8^+/E1114X^ compared to CHD8^+/+^-derived CblOs for the EN cell cluster (**Figure 4A**). To test whether these transcriptional alterations would impact electrophysiological properties, we performed MEA recordings at 130 days of differentiation (**Figure 4B)**. We observed a reduction in total network activity in CHD8^+/E1114X^-CblOs compared to CHD8^+/+^ ones (**Figure 4C**), whereas the activity occurring between consecutive network events indicated by MFR extraburst, was significantly increased in the mutant compared to controls (**Figure 4D**). These functional effects observed in CblOs from CHD8^+/E1114X^ at late maturation stages are compatible with an abnormal maturation of RL/oligodendrocytes, since both these cell lineages take part in the development of a proper neuronal network. These findings are also consistent with the virtual absence of INs in CHD8 mutants at 21 days of differentiation (**Figure 3D**). Immunostaining paired with tissue clearing aiming at identifying the Purkinje cells (PCs) marker CALB1, performed on CblOs at 130 days of differentiation, shows immunopositive thin cell bodies and elongated processes spreading around the surface of CblOs from CHD8^+/+^, while CALB1-positive cells in CHD8^+/E1114X^-derived CblOs exhibit an undifferentiated and rounded cell soma without ramified protrusions (**Figure 4E**)). Taken together, these results, in line with the excitatory/inhibitory neuron imbalance hypothesized for ASD patients^57,58^, point at an anticipated maturation of ENs at early differentiation stages, paired with the lack of properly mature inhibitory cells, ultimately leading to functional alterations of the neuronal network in CHD8^+/E1114X^ -derived CblOs. Additionally, the effects described for this CHD8 mutant in CblOs point to an opposite effect in the trajectory of cortical *vs.* cerebellar INs development according with our results obtained from unpatterned brain organoids^38^.

**FIGURE 4.**
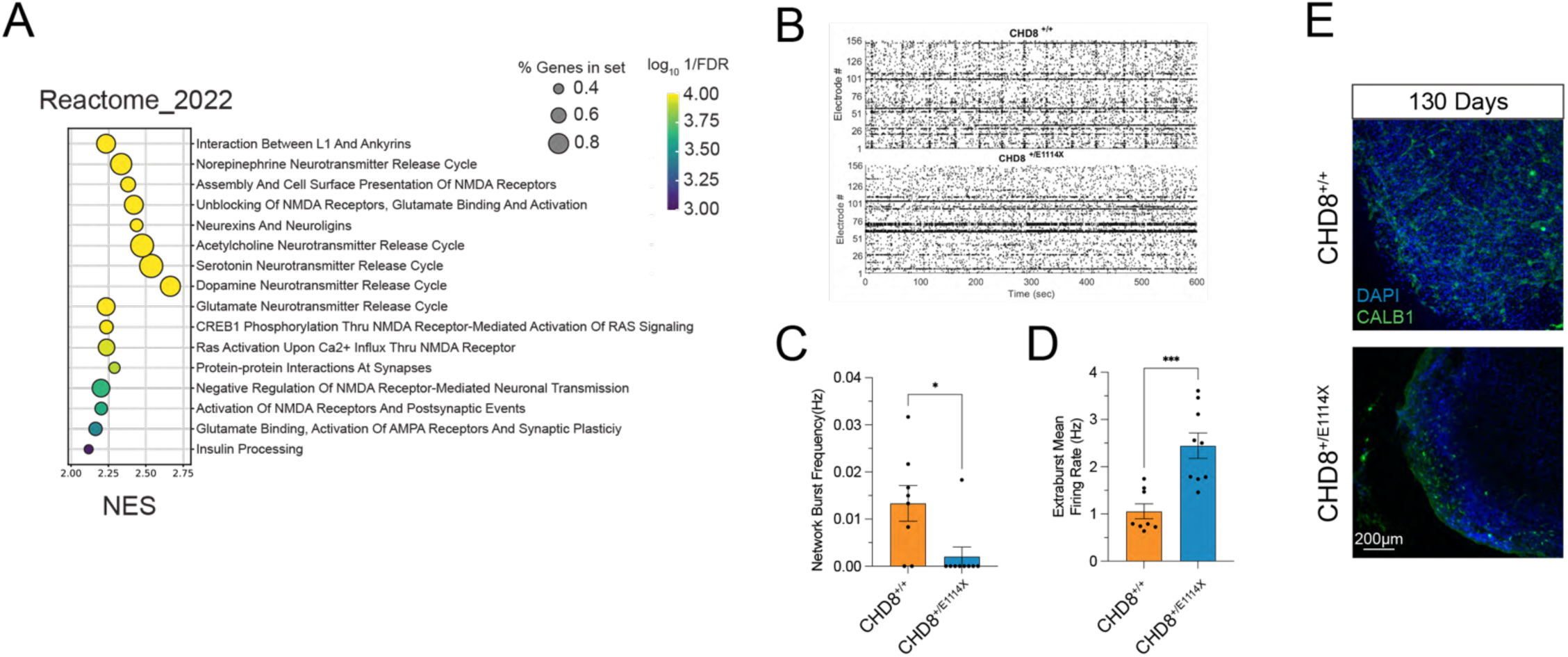
CHD8+/E1114X -derived CblOs show functional alterations of network activity at late developmental stages. (**A**) Functional enrichment analysis on Reactome gene sets performed by GSEA. Differential expression analysis by Wilcoxon test was performed to compare CHD8^+/+^ and CHD8^+/E1114X^ for EN/iGN clusters and the genes ranked by fold-change. The plot shows the NES (normalized enrichment score) and significance (-log10 FDR) for the gene sets that were found to be significantly enriched among the DEGs (FDR < 0.001). **(B)** Representative raster plots from MEA recordings of CHD8^+/+^ (upper panel) and CHD8^+/E1114X^ (lower panel) at 130 days of differentiation. **(C& D)** Histograms showing data from CHD8^+/+^ (orange) and CHD8^+/E1114X^ (blue)–derived CblOs at 130 days of differentiation. Network burst frequency is represented in (C). Mann-Whitney p= 0,0165 (*); CHD8^+/+^ n = 8 CHD8^+/E1114X^ n = 9). In (D), histograms show extraburst mean firing rate at the same timepoint. Mann-Whitney p= 0,0079 (***), CHD8^+/+^ = 5; CHD8^+/E1114X^ n= 5. **(E)** Representative spinning disk confocal microscope images of CHD8^+/+^ and CHD8^+/E1114X^ 130 days-old CblOs immunostained for CALB1 (green) and DAPI (blue).

### CADPS2 perturbations affects CblOs development

Having leveraged the CblOs model to assess the impact of CHD8 mutation on cerebellar development, we then sought to extend its application to the context of human evolution. To select the relevant Homo sapiens-specific SNV and recapitulate the effects of archaic alleles in CblOs, we combined a catalog of high frequency derived variants in Homo sapiens^59^ with cerebellar cortex enhancer data available from PsychENCODE Consortium. In the search for best candidates, we prioritized single nucleotide variants within regulatory regions depleted of archaic (Arc) derived mutations (so-called “regulatory islands”^16,60^), as well as regions associated with signatures of positive selection^61^ and depleted of introgression events (“introgression deserts” ^16,62^). Expression level of genes associated with such variants (using GREAT software^63^) in single-cell gene expression data from human fetal cerebellum ^6,64^ was also evaluated for top candidates. This led us to identify a mutation predicted to impact CADPS2, a gene encoded in the autism susceptibility locus 1 (AUT1) within chromosome 7 (**Figure 5A**). The SNV is found in an enhancer sequence embedded within intron 2, where the ancestral allele G was replaced by an A, found in most contemporary humans.

**FIGURE 5.**
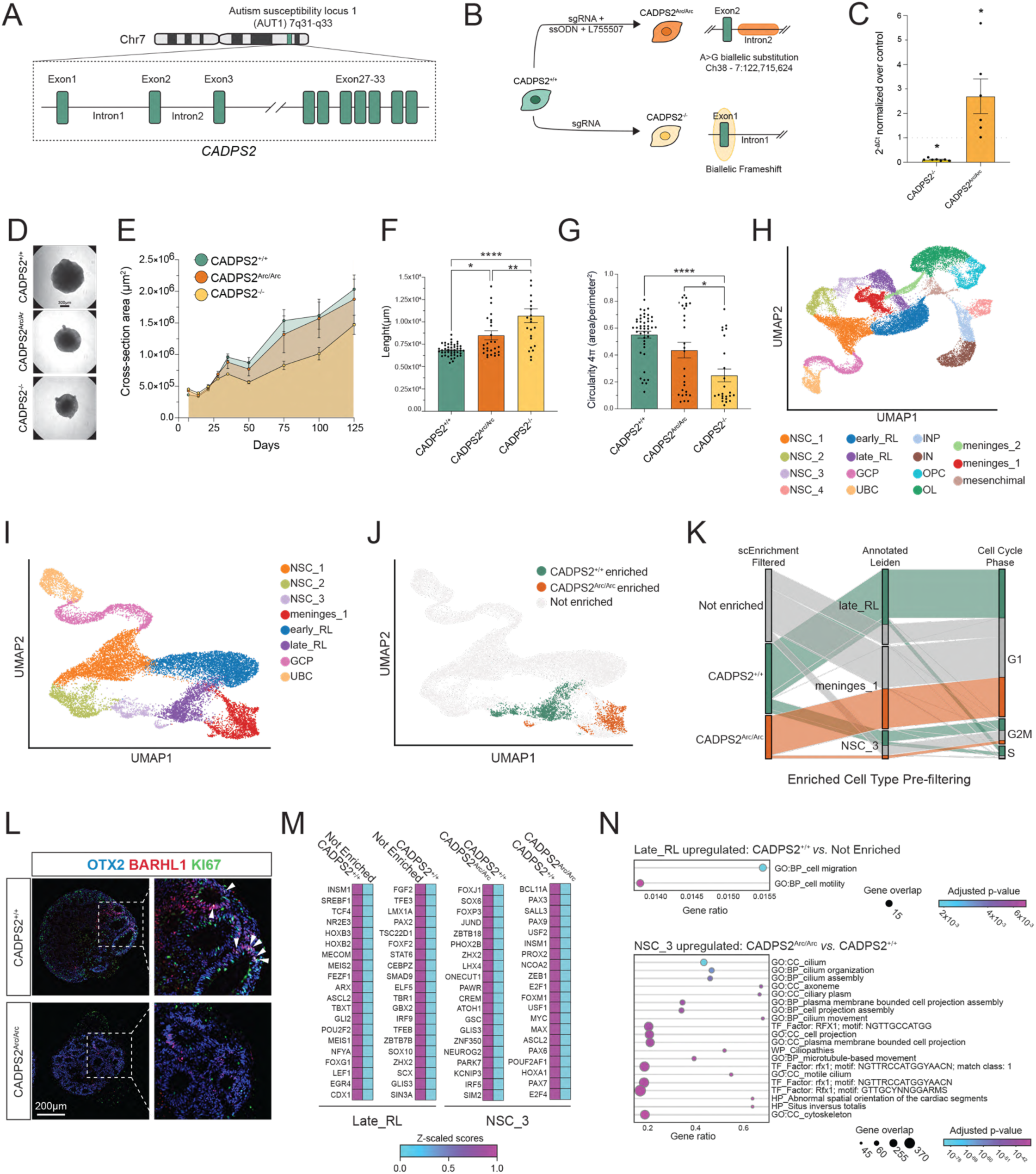
CADPS2 perturbations affect CblOs development. **(A)** Representative schema of the CADPS2 genomic structure localized within the autism susceptibility locus 1 (AUT1) in chromosome 7. **(B)** Schema describing the CRISPR/Cas9 approaches used to substitute within the second intron of CADPS2 (Ch38 - 7:122,715,624), the A residue in with a G (CADPS2^Arc/Arc^), by using an approach exploiting a single strand oligonucleotide (ssODN), and the homology-directed repair (HDR) enhancer L755507, along with the generation of a CADPS2 knockout line (CADPS2^-/-^) **(C)** Histograms showing qPCR from CADPS2^-/-^ (light orange) and CADPS2^Arc/Arc^ (dark orange)-hiPSCs. CADPS2^+/+^ (n=7) 0,4659±0,2527; CADPS2^-/-^ (n=7) 3,728±0,2157 Wilcoxon matched-pairs signed rank test: CADPS2+/+ *vs.* CADPS2-/- p= 0,0156 (*) CADPS2^+/+^ (n=6) 1,179±0,1627; CADPS2^Arc/Arc^ (n=6) -0,01097±0,4909. Wilcoxon matched-pairs signed rank test: CADPS2^+/+^ *vs.* CADPS2^Arc/Arc^ p= 0,0312 (*) **(D)** Representative transmitted light images from CADPS2^+/+^, CADPS2^-/-^ and CADPS2^Arc/Arc^ -derived CblOs at 35 days of differentiation. **(E)** Time course showing CblOs growth curves from 7 to 125 days of differentiation from CADPS2^+/+^, CADPS2^Arc/Arc^ and CADPS2^-/-^-derived CblOs. Measurements from four differentiation batches. **(F and G)** Histograms showing perimeter (**F**) and circularity **(G)** calculated from CADPS2^+/+^ (green), CADPS2^-/-^ (light orange) and CADPS2^Arc/Arc^ (dark orange)–derived CblOs at 125 days of differentiation from 4 rounds of differentiation. Circularity: CADPS2^+/+^ (n=45) 0,5516± 0,02562; CADPS2^Arc/Arc^ (n=28) 0,4363±0,05792 CADPS2-/- p=<0.0001 (****). One way ANOVA for multiple comparison was used: CADPS2^+/+^*vs.* CADPS2^Arc/Arc^ p=0,1045 (ns); CADPS2^+/+^ *vs.* CADPS2^-/-^ (n=24) 0,2485± 0,04774 CADPS2^Arc/Arc^ *vs.* CADPS2-/- p= 0,0128 (*). Perimeter: CADPS2^+/+^ (n=45) 6858 ±97,70; CADPS2^Arc/Arc^ (n=28) 8499± 504,3; CADPS2^-/-^ (n=24) 10692±771,3. One way ANOVA for multiple comparison was used: CADPS2^+/+^ vs. CADPS2^Arc/Arc^ p = 0,0149 (*); CADPS2^+/+^ vs. CADPS2-/- p= <0.0001 (****); CADPS2^Arc/Arc^ vs. CADPS2-/- p= 0,0040 (**). **(H)** UMAP embeddings of scRNAseq data from CblOs at 25 days colored by annotated clusters. Cells were clustered based on their expression pattern using the Leiden algorithm and the annotation of different cell types was generated on the base of published literature on cerebellar development. Unipolar brush cells (UBC), inhibithory neuron progenitors (INP) result from two independent differentiation rounds. **(I)** Integrated UMAP embeddings of CADPS2-expressing cells and their closest derivatives, colored coherently with the annotated clusters from (H). **(J)** UMAP embeddings of (I), colored by cells neighborhood enrichment status upon differential abundance between CADPS2^+/+^ (green neighborhoods) and CADPS2^Arc/Arc^ (dark organge neighborhoods). Non-enriched neighborhoods are colored in light-grey. **(K)** Sankey diagram showing the mapping for differentially abundant cell types (star methods) in CADPS2^+/+^ and CADPS2^Arc/Arc^ enriched neighborhood to annotated clusters and cell cycle phases. **(L)** Representative spinning disk confocal microscope images of CblOs from CADPS2^+/+^ and CADPS2^Arc/Arc^ at 25 days of differentiation. CblOs were stained for OTX2 (blue), BARHL1 (red) and KI67 (green). Harrow heads show the enrichment of triple-positive cells around ventricles. **(M)** TF activity analysis performed on the differential abundance clustering between CADPS2^+/+^ enriched *vs.* CADPS2^Arc/Arc^ enriched in NSC_3 and CADPS2^+/+^ enriched *vs.* not enriched domains and late_RL (see star Methods). **(N)** Functional enrichment of DEGs deriving from CADPS2^+/+^ enriched *vs.* not enriched domains (late_RL cell type), and from CADPS2^Arc/Arc^ enriched *vs.* CADPS2^+/+^ enriched domains (NSC_3).

To model the effect of this variant on cerebellar development, we exploited CRISPR/Cas9 to edit the selected locus in the control hiPSC line CTL04E already used for CblOs longitudinal characterization (thereafter referred to as CADPS2^+/+^). To avoid the introduction of additional SNV in the surrounding of this locus, we designed a sgRNA carrying a protospacer adjacent motif (PAM) identical to the genomic sequence. In addition, to generate a CADPS2^Arc/Arc^-line, we used the homologous direct recombination (HDR) enhancer L755507 along with a single strand ODN (ssODN) designed to flank the mutation’s region as described from Yu and colleagues in 2015^6,65^. To help with interpreting the effect of the SNV on directionality of expression, we also generated an additional line, starting from the same genetic background, introducing a frameshift mutation within the first exon of CADPS2 to generate a CADPS2^-/-^-knockout line and abrogate its expression. The CRISPR/Cas9 editing strategies and the oligonucleotides sequences used are illustrated in **Figure 5B** and **Supplementary Table S1** respectively. A detailed descriptions of all the quality checks performed on such lines can be found in methods. qPCR showed an abrogation of CADPS2 mRNA levels in CADPS2^-/-^ hiPSCs, whereas CADPS2^Arc/Arc^ showed a notable increase of CADSP2 levels in comparison to controls (**Figure 5C**). Similarly to what we did in the context of CHD8 mutations, we first assessed if changes in CADPS2 expression had an impact on CblOs development by measuring organoid growth over time. Although CblOs from both mutant lines showed variation in size and shape, the area under the curve calculated on cross-section areas measured from 7 to 125 days of differentiation was not significantly different compared to controls (**Figure 5D, E and Figure S5A**). However, other differences turned out to be significant: CblOs from CADPS2^-/-^ and CADPS2^Arc/Arc^ showed an increase in their perimeter length compared to controls, while circularity in CADPS2^-/-^ was reduced with respect of both CADPS2^+/+^ and CADPS2^Arc/Arc^ (**Figure 5F and G**).

In line with our results pointing to a preferential expression of NDD genes at early stages of CblO development, and the established association of CADPS2 with ASD and ADHD^34,35^, we performed scRNAseq with the three lines on CblOs at 25 days of differentiation obtaining 15 different cell clusters (**Figure 5H**). Most of the identified cell types were coherent in their transcriptional identity with cells observed in **Figure 3B** and consistent among genotypes and experimental replicates (**Figure S5B and S5C**). There was no cell-type specific enrichment for CADPS2^-/-^ distinct from the CADPS2^Arc/Arc^ one, so we focus on the latter in what follows. The marked expression differences of CADSP2 across conditions in the RL-derived lineage led us to isolate cell types expressing CADPS2 in that region (NSC_1, 2 and 3, early and late RL, GCPs, UBCs and meninges_1, **Figure 5I** and **Figure S5D**).

Based on their genotype, we identified a differential enrichment for CADPS2^+/+^ and CADPS2^Arc/Arc^ in CblOs (**Figure 5J**). The late_RL cluster was enriched in CADPS2^+/+^- cells, while meninges_1 were mostly composed of CADPS2^Arc/Arc^. In addition, both genotypes shared an enrichment within the NSC_3 cluster, but CADPS2^+/+^ and CADPS2^Arc/Arc^ occupied regions of the UMAP neatly clustered apart from each other. From imputation on the cell cycle phase obtained *via* transcriptomic data on each of these clusters (**Figure S5E**), we identified for CADPS2^+/+^ an enrichment of NSC_3 in both S and G2M phases whereas CADPS2^Arc/Arc^ cells are represented only in a limited fraction of cells in G2M phase (**Figure 5K)**. Controls were also enriched in S phase compared to archaic mutants in late_RL cells, and this finding was corroborated by immunostaining showing cells triple positive for OTX2/BARHL/KI67 and enriched in CADPS2^+/+^ (**Figure 5L**). In addition, most cells in G1 phase belonging to the control and CADPS2^Arc/Arc^ are enriched in late_RL and meninges_1 respectively, indicating the acquisition of a preferential genotype-based cell type identity.

We then computed the TF activity differences to characterize for each enriched genotype the molecular drivers resulting in specific cell type features (**Figure 5M**). This analysis highlighted for CADPS2^Arc/Arc^ cells within the NSC_3 cluster, the occurrence of TF related with the maintenance of sonic hedgehog (SHH) signaling, namely FOXJ1 and ATOH1. FOXJ1, a master regulator involved in ciliogenesis, has been associated with the acquisition of ependymal identity retaining neurogenic potential in brain cells^66^, whereas ATOH1 regulates proliferation of GCPs in the cerebellum *via* SHH signaling at the primary cilium^67^ in the external granule layer (eGL). These results are coherent with the functional analysis performed on DEGs calculated comparing the two genotypes within same cluster (**Figure 5N**), with CADPS2^Arc/Arc^-enriched cells showing an upregulation in cilia-associated terms in accordance with its higher FOXJ1 activity (**Figure S5F-S5G**). CADPS2^+/+^ enriched cells in the NSC_3 cluster instead, are associated with an increased activity of PAX6 and ZEB1. In the late RL cluster, we observed activation of TCF4, a TF associated with GCPs migration^68^ and related to the WNT/β-catenin pathway cell migration and motility are GO terms enriched in the late_RL cluster.

### O_2_ tension regulates CADPS2 expression

scRNAseq data from early developing CblOs, brought us to investigate how the ancestralizing SNV could affect fate acquisition. Specifically, we reasoned that an altered TF activity in CADPS2 ^+/+^ *vs* CADPS2^Arc/Arc^ lines could be at play.

We first resorted to ChIP-seq-derived position weight matrices from the JASPAR2024 database to predict in silico putative differences in TF binding at the relevant enhancer region (**Figure 6A**). In a contour of 37 nucleotides centered on the target A-G transition, we identified predicted binding sites for a total of 27 TFs (**Figure S6A).** Among those 27, CADPS2^Arc/Arc^ exclusively exhibited two consensus sequences for TCF7L2, a SFARI gene encoding TF factor involved in the β-catenin pathway^69^ and associated with ASD^70^, along with STAT1, a TF involved in NSCs self-renewal ^71^, also implicated in brain meninges cells proliferation^72^. CADPS2^+/+^ instead, exhibited 3 unique consensus sequences for ZKSCAN1, MYB and HIF1A. Among those, we focused our investigation on HIF1A, since it is expressed at high levels in GCPs ^73^ and has an established role in cerebellar development experimentally validated in mice ^36^. In particular, Kullmann and colleagues in 2020^31^, demonstrated that the hypoxic state of the developing cerebellum boosts Hif1a expression (modulating Zeb1 and Itgb1) maintaining granule cell progenitor proliferating progenitors in a proliferative state until vascularization creates normoxic conditions, at which point Pard6a stimulates granule cells to cease proliferation and begin migration.

**FIGURE 6.**
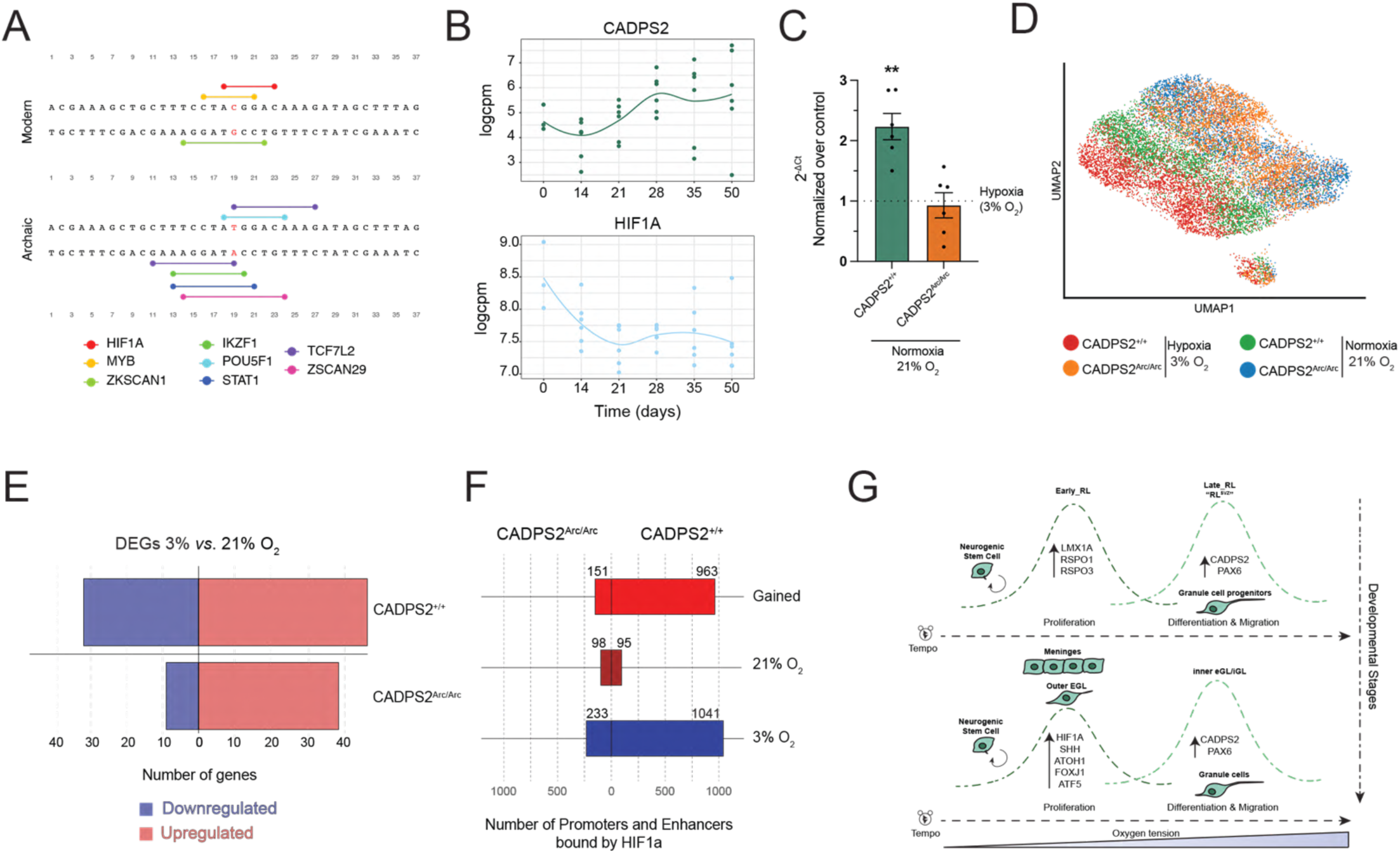
O2 tension regulates CADPS2 expression. **(A)** Representative schema showing the SNV surrounding region embedded in Intron 2 of CADPS2, along with the reconstruction of TF binding consensus sequences containing the SNV. The top strands correspond to the C/G sapiens-derived allele; bottom stands correspond to the T/A ancestral variant. Horizontal-dot capped sticks highlight the TF binding motifs. **(B)** Trend plots showing the longitudinal expression of CADPS2 (dark green) and HIF1A (light blue) from bulk RNAseq analysis. Data were obtained from three hiPSCs lines (CTL02A, CTL04E/CADPS2^+/+^; CTL08A) at 0, 14, 21, 28, 35, 50 days of differentiation, with two replicates per cell line. **(C)** Histograms showing the quantification of qPCR from CADPS2^+/+^ (green) and CADPS2^Arc/Arc^ (dark orange)-hiPSCs. Values were normalized on hypoxic conditions (3%O_2_=1). pValues were calculated on ΔCt: CADPS2^+/+^_3%O_2_ (n=6) 1,179± 0,1627; CADPS2^+/+^_21%O_2_ (n=6) 0,02060± 0,2463; CADPS2^Arc/Arc^ _3%O_2_ (n=6) -0,01097±0,4909; CADPS2^Arc/Arc^ _21%O_2_ (n=6) 0,3470±0,08005. Mann-Whitney with nonparametric test was used: CADPS2^+/+^_3%O_2_ *vs.* CADPS2^+/+^_21%O_2_; p=0,0043 (**); CADPS2^Arc/Arc^ _3%O_2_ *vs.* CADPS2^Arc/Arc^ _21%O_2_; p>0.9999 (ns). **(D)** UMAP embeddings of scRNAseq data from hiPSCs from CADPS2^+/+^ and CADPS2^Arc/Arc^ at the indicated O_2_%. **(E)** Histograms showing upregulated and downregulated genes at 21% O_2_ in comparison to 3% O_2_ conditions from scRNAseq of CADPS2^+/+^ and CADPS2^Arc/Arc^ hiPSCs. **(F)** Histograms from ChIP experiments showing the number of promoters and enhancer sequences bound by HIF1A at 3% (blue) and 21% (dark red) O_2_ in CADPS2^+/+^ and CADPS2^Arc/Arc^ hiPSCs and gained sequences bound by HIF1A (red) obtained comparing the two O_2_ conditions. **(G)** Schema recapitulating the developmental dynamics of rhombic lip derivatives and their relationship with O_2_ levels in the cerebellum associating cell types to the expression of specific genes, paralleling the situation obtained for the external granule layer (eGL) and inner granule layer (iGL).

Interestingly, HIF1A longitudinal expression in CblOs from three hiPSCs control lines revealed a trend symmetrically opposite to CADPS2, especially from 21 days of developments onward with a drop in HIF1A and a rise in CADPS2 expression (**Figure 6B**). Likewise, ITGB1 and ZEB1 peak when HIF1A is high, whereas PARD6A mimics CADPS2 expression (Figure S6B).

To test if indeed CADPS2 mRNA levels were sensitive to O_2_ tension in CADPS2^+/+^ compared to CADPS2^Arc/Arc^, we performed qPCR in hiPSCs. We observed that CADPS2 mRNA levels didn’t vary according to different O_2_ conditions in ancestralized lines, whereas in control hiPSCs normoxia (21% O_2_) causes a two-fold increase in the expression of the gene compared to hypoxic conditions (3% O_2_; **Figure 6C**). In light of this result, we tested if major transcriptional changes would have occurred in CADPS2^+/+^ and CADPS2^Arc/Arc^ lines at different O_2_ levels. To do so, we performed scRNAseq in hiPSCs from both genotypes maintained at either 3% or 21% O_2_ (**Figure 6D**). CADPS2^+/+^ and CADPS2^Arc/Arc^ UMAP showed a divergent transcriptomic profile, with controls displaying an increased number of DEGs in comparison with their ancestralized counterpart, indicating a generalized susceptibility of cells to O_2_ tension (**Figure 6E**). This finding was further corroborated by a ChIP assay targeting HIF1A, performed in the same experimental settings (**Figure 6F**). Results of this experiment showed that CADPS2^+/+^ hiPSCs in hypoxia have an increased number of promoter and enhancer sequences bound by HIF1A in comparison to CADPS2^Arc/Arc^ cells. The number of sequences bound by this TF dramatically drops at 21% O_2_, when the number of HIF1A-bound promoter and enhancer sequences became equally represented in both genotypes.

The CADPS2^+/+^ sensitivity to O_2_ levels, tied to HIF1A regulation, led us to the conclusion that in Homo sapiens CADPS2 expression levels are high in those regions that are well oxygenated, such as the late RL, plausibly corresponding to the richly vascularized human specific outer RL (RL^VZ^)^74^, and low in the outer eGL, adjacent to the meninges, where HIF1A expression is high (and where as we saw above CADPS2^+/+^ is not enriched). In consonance with Kullmann et al. 2020^36^, CADPS2 expression is expected to rise as GCPs migrate away from the outer eGL and into the inner granule layer (iGL) reflecting the early/late RL situation, whether HIF1A binding essentially delays the rise in CADPS2 expression and granule cell maturation. (**Figure 6G**). More generally, our work supports the claim that meninges play a critical role in cerebellar development^75–78^, and possibly beyond^79–81^.

## DISCUSSION

Our high-resolution benchmarking establishes CblOs as modelling resource for studying the impact on human cerebellar development of genetic variants associated with contemporary neurodiversity and its most recent evolutionary underpinnings. The early stages of cerebellar development as captured in CblOs derived from three control hiPSC lines, reveal a highly dynamic changes in the expression of a specific gene core enriched for variants causally associated to cardinal neurodevelopmental conditions. This provides the foundation for probing how these variants bias divergent neurodevelopmental trajectories that eventually underlie distinct cognitive/behavioral traits among contemporary humans and also, ceteris paribus, across closely related species. This density of such pathophysiologically relevant expression changes in early development extends to cerebellar organoids the paradigm emerging from their cortical counterpart, on how earliest development captures most of the mutation-specific transcriptional changes that are then propagated along development in convergent modules^82^.

The possibility to study through CblOs the early impact of such changes *in vitro*, allowed us to describe cellular trajectories and functional consequences of mutations in the high penetrance and ASD-related gene CHD8. We identify for this gene an effect on the developmental tempo of RL-derived cells, INs and OLs, that synergically reverberate through CblOs maturation hampering the functional electrophysiological properties of mature organoids. Those results highlight the ability of CblOs to provide over time, a comprehensive view of cell types often investigated into isolated contexts as the effect of CHD8 was studied in the RL lineage^39,83^ or glial cells^84–86^.

Our work also shows that the same stage at which of CblOs differentiation captures early cerebellar effect associated to neurodevelopmental disorders, corresponds to a sensitive period in which CblOs can capture the evolutive divergence between Sapiens *vs*. Neanderthal and Denisovans. The study of archaic variants associated with CADPS2, showed a distinct patterning in the acquisition of a RL-derived and meningeal cell fate lending credence to the idea that significant differences in brain growth trajectory implicates the interplay of brain tissues with structures like the meninges and ventricles^87,88^. In this light, the shape of the neurocranium may be the outcome of a set of coordinated changes across brain, overlying meninges (fibroblasts, vasculature and immune cells), cranial base^89^ and calvarium with non-strictly brain structures determining the direction of brain expansion during development and affecting endocranial shape, as is also clear in the case of diseases such as hydrocephalus, Dandy-Walker syndrome, Joubert syndrome and other ciliopathies.

Additionally, our findings on CADPS2 link O_2_ tension to the maturation of the RL lineage, extending the idea of delayed neuronal maturation (‘neoteny’) to the cerebellum, contrary to previous claims characterizing this effect as cortex-specific ^90^. It will be worth exploring in the future if the much more extensively discussed Purkinje cell-related deficits in conditions like ASD, but also evolutionary modifications, may also be rooted in very early granule cell perturbations provoking a neotenic cascade: delaying the maturation of granule cells and thus also the later-maturing Purkinje neurons. Relatedly, our ASD-modelling in cerebellar organoids points to the need to broaden our focus beyond the Purkinje layer, as we revealed the critical role of the RL in altering developmental trajectories. The involvement of RL-derived cells in fact, has been recently described in the context of knockout mice for the ASD gene Shank3^91^ and it will be worth examining in the future how far among ASD-related candidate this extends.

It will also be worth combining organoid modelling of distinct brain regions, possibly with assembloids^20,59^, to examine in more detail the reported cerebellum-specific volume reduction in patient lines that contrasts with the overall macrocephalic profile reported, pointing to region-specific asynchronies that may contribute to dysconnectivity profiles.

For ASD and related neurodevelopmental disorders, it will be important to generate organoids for many more candidate mutations impacting different genes to determine the range of neurodivergence. In a similar vein, on the evolutionary side, it is important to remember that no single mutation is sufficient to capture a complex phenotype ^59,92,93^, and we hope that our study can serve as a resource to explore additional differences for genes that are also known to have a divergent profile and that belong to different clusters in our analysis (e.g., GLI3, TCF4, OTX1, STAT2).

## LEAD CONTACT

Requests for information, resources and reagents should be sent to Prof. Giuseppe Testa (<u>Giuseppe.testa@fhtorg</u>).

## MATERIALS AVAILABILITY

Plasmids and cell lines generated in this study are available upon request to the lead contact with a completed materials transfer agreement.

## DATA AND CODE AVAILABILITY

The lead contact will provide, upon justified inquiry, any supplementary details needed for further examination of the study’s findings.

## ACKNOWLEDGMENTS

This work received the institutional support by the Coordinated Research Centre on Organoid Biology (University of Milan), the European Union’s Horizon 2020 research and innovation program grants R2D2-MH (101057385 to G.T.), the European Union’s Horizon 2020 research and innovation program grants RE-MEND (101057604 to G.T.), the BBRF YI grant 2023 (31998 to D.A.), and the Telethon foundation grant n°GGP19295. O.L. enrolment as PhD student in System Medicine was possible thanks to the European School of Molecular Medicine (SEMM). We thank Eugenia Ricciardelli along with Paolo Ferrari for their support on transcriptomic experiments, and Christopher Schmied for helping in designing the macro for immunostaining quantification. We want to thank J.L. Rubenstein for having shared with our group the Lenti-Dlxi1/2b::eGFP construct and express our gratitude to Prof. Kimberly A. Aldinger and Prof. Kathleen J. Millen for sharing the transcriptomic data on human fetal cerebellum. We are grateful to Prof. Gaia Novarino for sharing the CHD8 lines. Funding bodies did not have any influence on study design, results, and data interpretation or final manuscript.

## AUTHOR CONTRIBUTIONS

G.T. and C.B. conceived and supervised the work; G.T., C.B. and D.A. conceptualized and designed experiments; DA performed most of the experiments. C.C., M.B., A.T. performed the bioinformatic analysis related to CHD8. D.C., A.Vitriolo and J.M. performed the bioinformatic analysis related to CADPS2 section. O.L. contributed to the bioinformatic analysis on both the CHD8 and CADPS2 section, also providing support for selected experiments. A.P. performed experiments related to CHD8 A.Valente performed experiments on CADPS2 lines. L.C. helped in cell culture & image analysis. F.M. performed MEA recordings and analysis.D.A., C.B and G.T. wrote the manuscript with contributions from all the authors that read and approved the final version of this work.

## DECLARATION OF INTERESTS

The authors declare no competing interests.

## Materials & Methods

### Key resources table

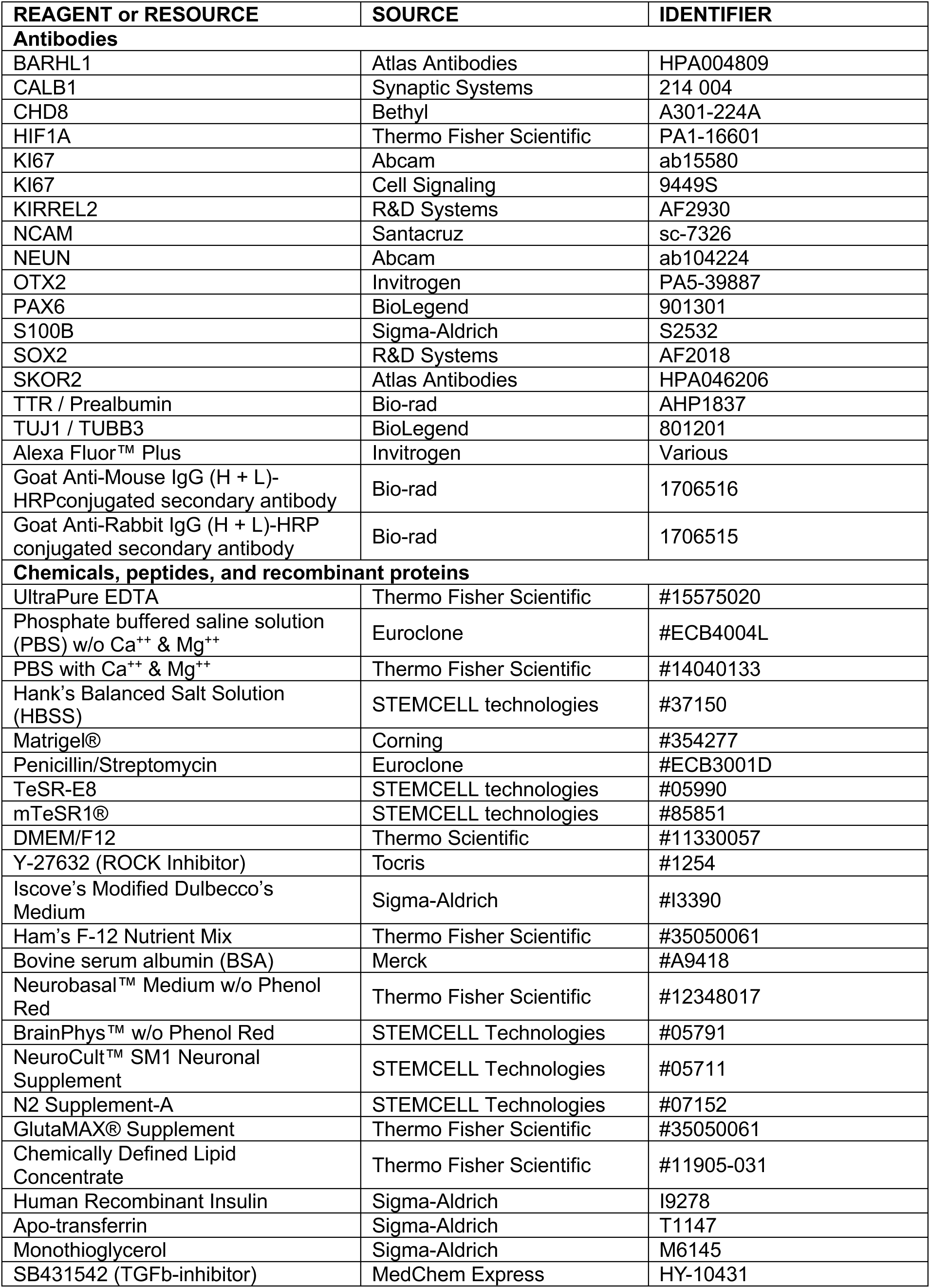
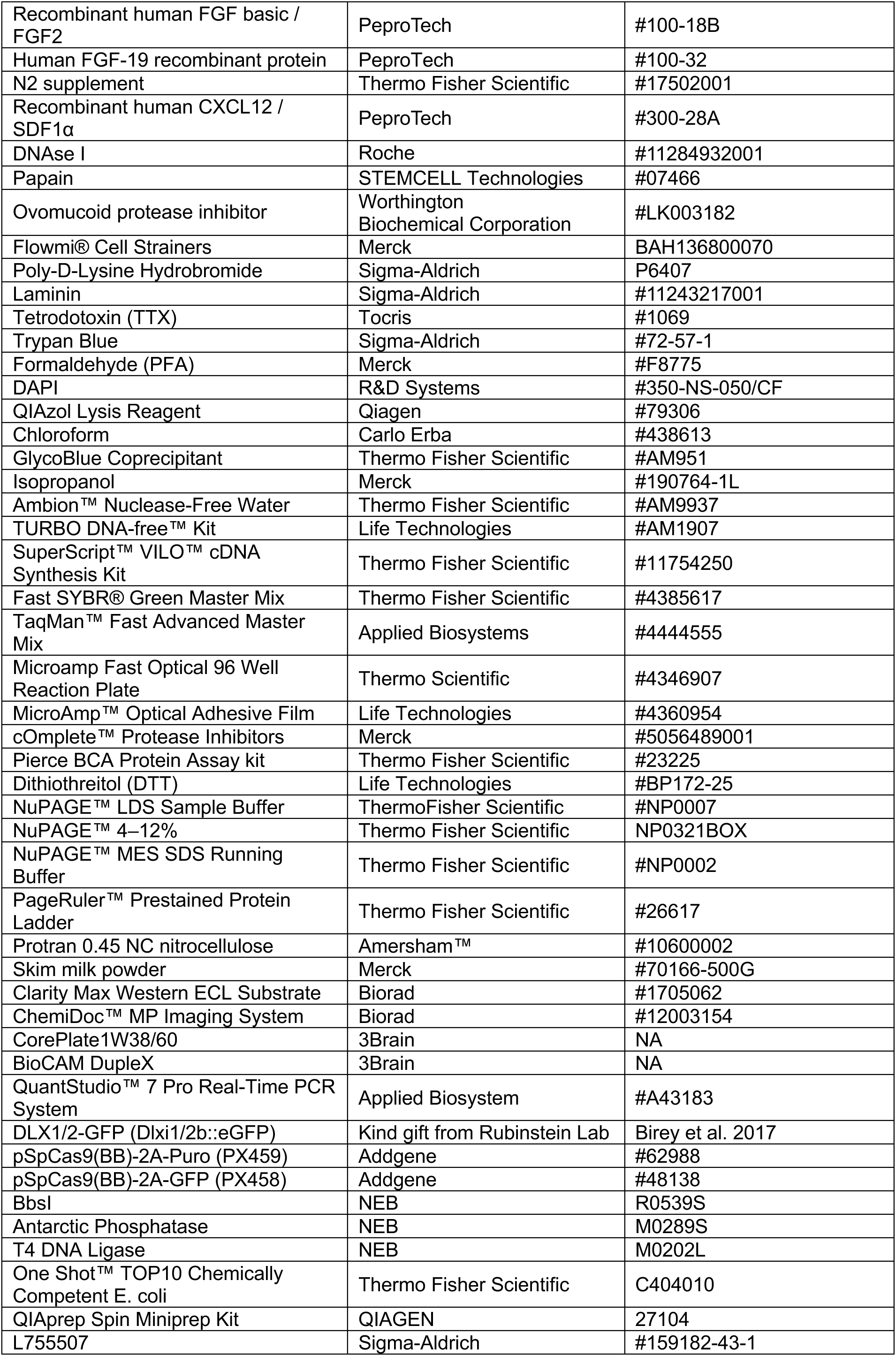
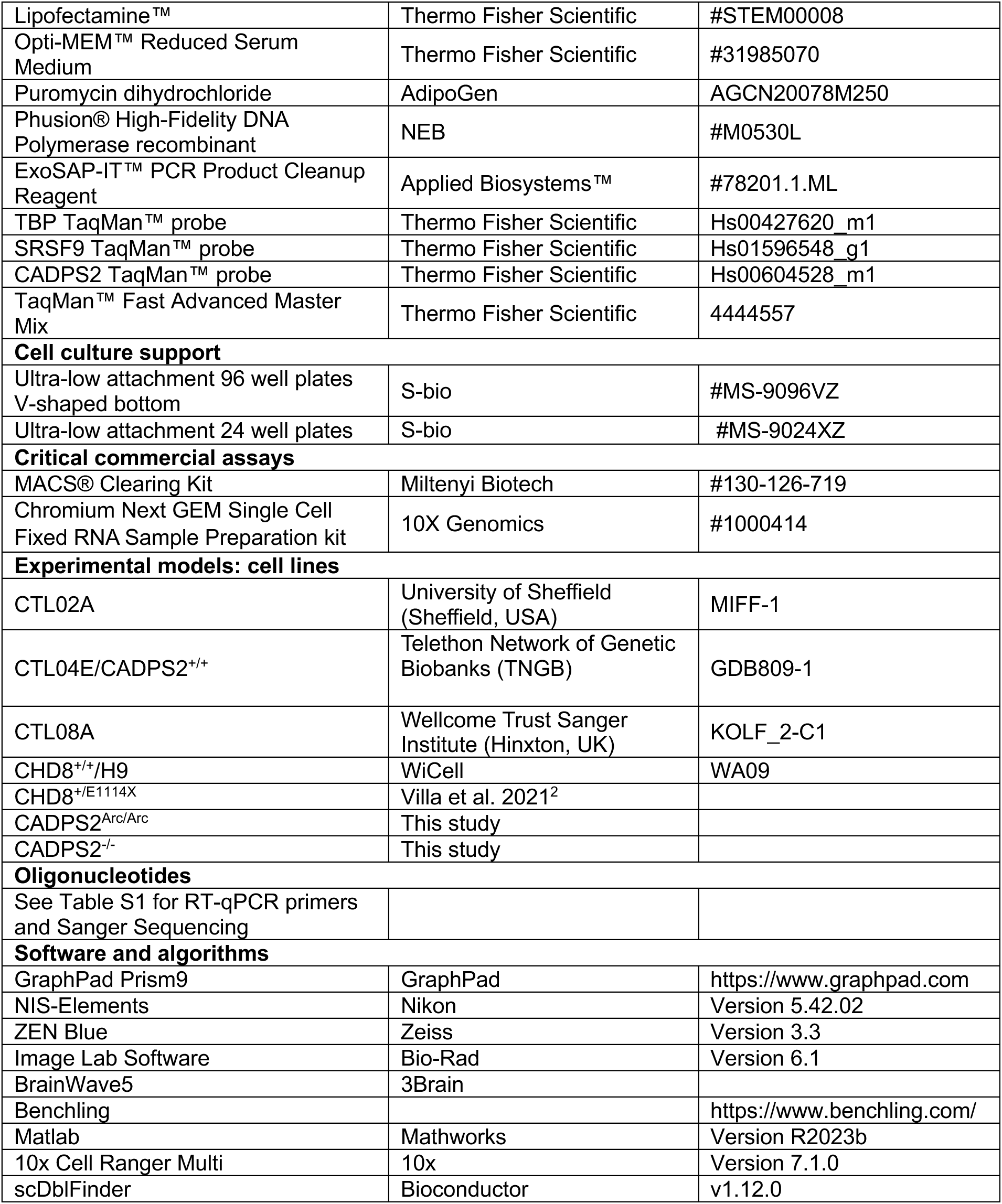

### EXPERIMENTAL MODELS

#### hiPSCs & hESCs maintenance

hiPSCs (CTL02A, CTL04E and CTL08A) were cultured at 37C°, 5% CO_2_ and 3% O_2_ in TeSR-E8 supplemented with penicillin/streptomycin(100u/ml). hESCs (Feeder-independent human H9 ES cells/CHD8^+/+^) and their derivatives CHD8^+/E1114X^ (Villa et al. 2021)^38^, were cultured in mTeSR1® medium supplemented with penicillin/streptomycin (100u/ml) at 37C°, 5% CO_2_ and 20% O_2_. Cells were grown on dishes coated with Matrigel® diluted in DMEM/F12, with daily media change. Once cells reached the 60-80% of confluency within the plate, were propagated by detachment of isolated colonies with 0.5 mM UltraPure EDTA diluted in phosphate buffered saline solution.

#### Cerebellar Organoids differentiation

Cerebellar organoids were generated using an adapted version of the protocol described in Muguruma et al. 2015^24^. Briefly, cultures at 80% confluency were single-cell dissociated, with UltraPure EDTA 0.5mM at 37C° for 15min, counted with a Countess 3 Automated Cell Counter (Thermo Fisher) in Tripan Blue, resuspended at the final concentration of 40’000 cells/ml in mTeSR1 complemented with 10μM ROCK inhibitor (RI) and seeded in ultra-low attachment 96 multiwell plates V-shaped bottom at the density of 6000 cells/150 ml. Embryoid bodies (EBs) formation was induced by centrifuging the plates at 200 g for 1 minute. Media change was performed as follows: after two days, growth factors complemented differentiation medium (gfCDM) composed by Iscove’s Modified Dulbecco’s Medium and Ham’s F-12 Nutrient Mix 1:1, GlutaMAX® Supplement 0.5%, BSA 5mg/ml, Chemically Defined Lipid Concentrate 1%, Apo-transferrin 15 μg/ml, Human Recombinant Insulin 7μg/ml, Monothioglycerol 450μM, penicillin/streptomycin 100U/ml, was added along with 10μM RI, 10μM TGFb-inhibitor (SB431542) and 50 ng/ml FGF2 supplementation. gfCDM supplemented with RI, TGFb-inhibitor and FGF2 was replaced again on day 7. On day 14, gfCDM with 100 ng/ml of FGF19 was used. From day 21 onward, gfCDM was replaced by Neurobasal™ Medium, 1% GlutaMAX®, 1% N2 supplement, 100 U/ml penicillin/streptomycin (Neurobasal + N2). SDF1α 300ng/ml was added to the media at day 28. On day 35, upon CblOs transfer into 24mw ultra-low attachment, weekly media changes with Neurobasal + N2 were performed. hiPSCs-derived CblOs were grown at 3% O_2_ until 21 days and moved at 21% O_2_ at 21 days, while CblOs derived from hESCs were kept at 21% O_2_ since day 0.

### METHODS DETAILS

#### Growth Curves

Transmitted-light images of 10-12 independent CblOs *per* condition from 7 to 125 days of differentiation were acquired by using an AxioVert.A1 microscope (Zeiss), equipped with a Plan-Neofluar 5X/0,16 or Plan-Neofluar 10X/0,3 dry objective. Semi-automatic detection of the major cross-section areas of the cerebellar organoids was obtained by applying an automatic intensity threshold based on the Otsu algorithm. For images with a low signal-to-noise ratio, the threshold was adjusted by a numeric modifier that was visually evaluated.

#### Infection and live imaging

50% confluent hiPSCs colonies were infected with DLX1/2-GFP (Dlxi1/2b::eGFP), 1MOI. Once hiPSCs reached the 80% of confluency, CblOs were generated and maintained until 90 days of differentiation for image acquisition.

#### Immunostaining followed by tissue clearing

Immunostaining followed by tissue clearing was performed according to the manufacturer indications (Miltenyi Biotec). At least 4-5 organoids from a single differentiation round were used for each experimental condition. Samples were acquired with a Nikon Ti2 microscope equipped with CREST v3 spinning disk and a PlanApo λ 20X/0.75NA dry objective. All quantifications were performed in Fiji ImageJ, processed with Microsoft Excel, and plotted in GraphPad Prism. Images were analyzed blindly. BARHL1 and S100B occupancies were calculated as percentage of the total area occupied by BARHL1+DAPI or S100B+DAPI signals, respectively. For each z-stack, a single sub-stack was selected based on the quality of DAPI signal and rolling ball background subtraction method was applied. Images were segmented with automatic intensity threshold based on the MaxEntropy (for BARHL1) or Moments (for S100B and DAPI) algorithm. A particle size ranging from 5px*^2^* to 70 px*^2^* was considered.

#### Multielectrode array (MEA)

CblOs were attached on Poly-D-Lysine (0.1mg/ml)/Laminin (0.04mg/ml)-coated CorePlate1W38/60 MEA and organoids were maintained in Brainphys supplemented with N2 Supplement-A and NeuroCult™ SM1 Neuronal Supplement for 5-7 days before spontaneous activity recordings. To assess the *bona fide* dependence of electrical events from glutamatergic transmission, 1mM tetrodotoxin (TTX) was applied at the end of every recording for 5 additional minutes. Recordings were performed using a BioCAM DupleX and the BrainWave5 software. CblOs activity was recorded for 10 minutes, with a sampling rate of 10kHz. A Bandpass filter (300-3000Hz) was applied to the traces before performing spike detection using the Precise Timing Spike Detection (PTSD) algorithm of the BrainWave5 software^94,95^. The threshold for the spike detection was set to 8 standard deviations, 1ms of refractory time and 1.5ms peak lifetime. For subsequent analysis, only the channels with a mean firing rate greater than or equal to 0.2 spikes/sec were considered active. Network Burts (NBs) were identified as major population events involving at least 12% of active electrodes. The neural activity trace was segmented into 50ms time windows, and for each bin, the total number of spikes (NP) and the number of active channels (AC) were computed. Subsequently, both NP and AC time series were smoothed using a median filter of order 20. The filtered NP and AC values were then multiplied to generate a composite parameter referred to as Product. A NB was defined as any event in which the Product exceeded the grater of either the mean value plus three the standard deviations or an absolute threshold of 10.

#### RNA extraction

Samples were harvested and washed with PBS with Ca^++^ and Mg^++^. RNA extraction was performed by using QIAzol Lysis Reagent and Chloroform from 4-6 CblOs per condition. Pellets were visualized with GlycoBlue Coprecipitant and the RNA was precipitated with Isopropanol. Samples were incubated overnight at -80°C to let RNA precipitate. The following day, samples were centrifuged at 12’500 rpm for 15 minutes at 4°C and the supernatant was removed. RNA pellets were washed twice with 75% EtOH and the samples were re-centrifuged at 12’500 rpm for 15 minutes at 4°C. Upon ethanol removal, pellets were air-dried for 10-15 minutes at RT and resuspended in the appropriate amount of Ambion™ Nuclease-Free Water. Samples were stored at - 80°C. For bulk RNAseq, samples were additionally treated with TURBO DNA-free™ Kit.

#### qPCR

Reverse transcription of the extracted RNA from cells and organoids was performed using the SuperScript™ VILO™ cDNA Synthesis Kit as indicated by manufacturer’s instruction. Samples were amplified with Fast SYBR® Green Master Mix in Microamp Fast Optical 96 Well Reaction Plate, 0.1ml. Plate were sealed with a MicroAmp™ Optical Adhesive Film, then centrifuged briefly to spin down the samples and remove any air bubbles. Customized oligos used for qPCR are listed in Table S1, while to test CADPS2 expression, TaqMan™ Fast Advanced Master Mix was used. TaqMan™ probes are listed in the **Tab Materials.** Reactions were run on QuantStudio™ 7 Pro Real-Time PCR System.

#### hESCs immunostaining

60-70% confluent hESCs seeded on a 18mm diameter glass slide and cultured on a 6 multi-well plate were fixed in PFA 4%. Cells were permeabilized with PBS-0.1% Triton X-100 for 10 minutes, followed by blocking with PBS-10% Donkey serum for 30 minutes. Cells were incubated with primary antibodies overnight at 4C°, washed three times with PBS and incubated with secondary antibody for 45 minutes. Antibodies used are listed in **Tab Materials**. Samples were acquired with a Nikon Ti2 microscope equipped with CREST v3 spinning disk and a PlanApo λ 20X/0.75NA dry objective.

#### Colocalization analysis

Colocalization analysis was performed in Fiji. The nuclear area of cells was segmented based on a maximum intensity projection based on the DAPI channel applying an automatic intensity threshold based on the Huang algorithm. The projection was then filtered using a median filter with a radius of 5 pixels and the rolling ball background subtraction method with sliding paraboloid was applied. Cell area was segmented based on a maximum intensity projection of the measurement channel and a Gaussian filter with sigma = 10 pixels was applied to the projection. An empirically determined global background value was used for background subtraction and an automatic intensity-based threshold was applied using the triangle algorithm. CHD8 mean intensity was measured both on the area of segmented nuclei and the total cell area (excluding the nucleus) by using an ImageJ/Fiji batch macro. For intensity measurements, a sub-stack obtained averaging projection in a contour of 10 slices around the brightest *z*-stack. Mean intensity was then measured on both segmented nuclei and cell area.

#### Protein extraction and Western Blot

*<u>Cytoplasm/nuclear fractions</u>:* cells were resuspended in 3V Low Salt Buffer (100mM HEPES pH 6.8, 5mM KC, 5mM MgCl2, 0.5% NP40 and cOmplete™ Protease Inhibitors), incubated 10 minutes on ice and then centrifuged at 4°C, 2000 rpm for 3 minutes. Nuclei were resuspended in 2V of High Salt Buffer (100mM HEPES pH 6.8, 250 mM NaCl, 5mM KC, 5mM MgCl2, 0.5% NP40 and cOmplete™ Protease Inhibitors), incubated on a rotating wheel at 4°C for 30 minutes with occasional homogenization using a 25G needle. The samples were centrifuged at 4°C, 2000 rpm for 3 minutes and the supernatant kept as nuclear fraction. Proteins were quantified with the Pierce BCA Protein Assay kit. *<u>Whole-protein lysates</u>*: samples were washed with PBS with Ca^++^ and Mg^++^ and lysed in denaturing conditions with NaCl 150mM, EDTA 1mM, Tris HCl 50mM, 1% Triton 100X and 1% SDS 20X. cOmplete™ Protease Inhibitors were added fresh. Samples were sonicated for 30 seconds and proteins retrieved after lysates clarification by centrifugation at 4°C, 12000 rpm for 10 minutes. *<u>Western Blot</u>:* protein samples were supplemented with Dithiothreitol (DTT) 500 mM and NuPAGE™ LDS Sample Buffer, then boiled at 95°C for 3 minutes. Extracts were separated on NuPAGE™ 4–12% gels in MES SDS Running Buffer and the separation followed with PageRuler™ Pre-stained Protein Ladder. Proteins were blotted on Protran™ Nitrocellulose membrane (0.45um) in transfer solution (0.25 M Tris, 1.92 M glycine, 20% methanol and 0.03% SDS). Membranes were quenched for 30 minutes in 5% skim milk powder diluted in Tris-buffered saline (10 mM Tris, 150 mM NaCl) with 0,1% Triton 100X. The same buffer was used for primary and secondary antibody incubation. Bands were revealed after a Clarity™ Western ECL Substrate exposition and acquired with ChemiDoc MP Imaging System. Bands quantification was performed with Image Lab™.

#### Single cell RNAseq

*<u>Organoids dissociation and RNA</u>:* 6 to 8 CblOs at 21-25 days of differentiation for each experimental condition were transferred into wells of a 24-multiwell plate and washed with PBS w/o Ca^++^ and Mg^++^. PBS was then replaced with the dissociation solution (Papain 30 U/ml; DNaseI 125 U/ml in Hank’s Balanced Salt Solution) and organoids were kept in orbital shaking (120rpm) for 10 minutes at 37°C. Organoids were triturated 5/6 times at RT to obtain a suspension consisting primarily of cell aggregates. Cell aggregates were incubated for other 5 minutes at 37°C in shaking. Aggregates were again triturated 4/5 times to obtain a single-cell suspension. The enzymatic reaction was inhibited by adding ovomucoid protease inhibitor (10mg/ml) in HBSS and cells centrifuged at 300 g for 5 minutes. Cells were resuspended in 1 ml of PBS w/o Ca^++^ and Mg^++^ with 0.04% BSA and counted. Remaining cell aggregates were removed filtering the cell suspension through a 70μm Flowmi® Cell Strainers. Cells were fixed in Chromium Next GEM Single Cell Fixed RNA Sample Preparation kit according manufacturer’s protocol for library preparation.

#### Generation of CADPS2 transgenic lines

##### sgRNAs and ssODN design

sgRNA for CRISPR/Cas9 gene editing were designed based on the GRCh38.p7 genome assembly using the online tool Benchling. sgRNA for CADPS2^-/-^ were designed to target the first CADPS2 exon, while CADPS2^Arc/Arc^ sequences targeted the second intronic region of the gene. In both cases, the sequences were chosen based on their on-target and off-target score. To introduce the A to G transition we designed a 200bp-long ssODN carrying the SNV enclosed within the surrounding chromosomal intronic sequence. Phosphorylated sgRNAs and the ssOND were provided by ThermoFisher and oligonucleotides are listed in Table S1.

*<u>Generation of the sgRNA expression constructs</u>:* pSpCas9(BB)-2A-Puro plasmid were digested with the BbsI restriction enzyme and de-phosphorylated with Antarctic Phosphatase. The ligation between the de-phosphorylated linearized pSpCas9(BB) plasmid and the sgRNA insert was performed using the T4 DNA Ligase.

*<u>Bacteria transformation and Sanger-sequencing sample preparation</u>:* One Shot™ TOP10 Chemically Competent E. coli were heat-shocked with 25ng of ligation products and cultured in S.O.C. medium at 300rpm for 1 hour. Transformation product was seeded on a pre-warmed Luria Agar petri dishes in presence of Ampicillin and incubated overnight at 37°C. Single colonies were singularly picked and amplified. Plasmids were extracted with QIAprep Spin Miniprep Kit, quantified and sent to the Eurofins (https://www.eurofins.it/) for Sanger sequencing on the U6-promoter. *<u>Cell</u> <u>Lipofection</u>*: transgenic lines were generated starting from the hiPSC control line CTL04E (CADPS2^+/+^). 80-85% confluent cells were detached with Accutase® and counted. Only plates with 80-95% of cell viability were considered to proceed with the experiment. 2,5*10^5^cells/well were seeded in a Laminin pre-coated 6 multi-well plate and maintained in mTeSR1® with Y-27632 with daily media change. On the following day, cells were incubated with a mix of 5ml of Lipofectamine™ Stem Transfection Reagent and 1,25mg of DNA in Opti-MEM™ Reduced Serum Medium. CADPS2^Arc/Arc^ cells were generated adding to the transfection mix along with the sgRNA-expressing vector, 1,25mg of ssODN. The pSpCas9(BB)-2A-GFP (PX458) vector was parallelly used as transfection positive control. After 24hours, lipofection media was supplemented with 0,3mg/ml of puromycin and for the following two days media changes in presence of puromycin were performed daily. To generate CADPS2^Arc/Arc^ cells, 5mM of L755507 was added at every media change. Isolated clones resistant to puromycin were manually picked and replated. *<u>Clones screening</u>*: genomic DNA was extracted from isolated clones following the protocol by McManus Lab (https://mcmanuslab.ucsf.edu/protocol/dna-isolation-es-cells-96-well-plate). Regions of interest were then amplified through PCR with Phusion® High Fidelity DNA Polymerase following manufacturer’s instruction (Primers in Table S1), the PCR products purified with ExoSAP-IT™ PCR Product Cleanup and sent to Eurofins for sequencing. Genomic stability was assessed through CNV analysis. As in both CADPS2^Arc/Arc^ and CADPS2^-/-^ we observed a duplication occurred within the Chr. 20 of both lines and an additional duplication of Chd.17 in CADPS2^-/-^. We then intersected, the genes encoded within the CNV and scRNAseq gene expression results from CblOs at 25 days of differentiation, generating a gene regulatory network to exclude the possibility that such rearrangements would have an impact on the differentiation of CblOs.

### Chromatin immunoprecipitation followed by sequencing (ChIP-Seq)

ChIP-seq targeting HIF1A was performed in duplicate for CTL04E and CADPS2^Arc/Arc^ hiPSCs at both 3% and 2% O_2_, using 10x^7^ cells *per* sample according to the procedure described in Pereira et al. 2025^96^.

### Bulk RNAseq analysis

***CblOs benchmarking*** - For the protocol benchmarking, each of the three hiPSC lines was differentiated and profiled with bulk RNAseq at the following time points: hiPSC stage (Day0), Day14, Day21, Day28, Day35 and Day50. Excluding hiPSCs profiled once, sample belonging to this cohort (n=33), were obtained from two differentiation rounds of CblOs. *<u>Alignment</u>* RNAseq FASTQ data were quantified at the gene level using Salmon. GRCh38 Genecode v35 was used as reference for quantification and annotation. Data will be available in ArrayExpress public repository upon publication. Bulk RNASeq analyses were performed with R version 4.2.1 unless specified differently. *<u>Differential Expression Analysis</u> (*DEA) was performed in a stage-wise approach comparing each differentiation stage with the previous time-point using DESeq2^97^ v1.36.0. Protein coding genes were selected for differential expression analysis. Gene filtering was applied to discard not expressed and lowly expressed genes (10 reads in at least 5 samples for Day14 *vs.* hiPSCs; 20 reads in at least 6 samples for all the other comparisons). In addition to line identity, the information about QC performance was used as a covariate in the statistical model when appropriate (all comparisons but Day14 vs hiPSCs). Fold-change shrinkage was applied using ’ashr’ algorithm in DESeq2. For each comparison, the number of DEGs, split in upregulated or downregulated, was represented by bar plots. *<u>Functional Enrichment Analysis</u>* DEGs were selected imposing a more stringent threshold (FDR < 0.05 and absolute log2FC > 1). Functional enrichment analysis for the Biological Process domain of the Gene Ontology was performed by TopGO v2.48.0 for every comparison, dividing genes in up- and downregulated and setting the following parameters: ‘weight01’ as algorithm, ‘fisher’ as statistics and 15 as ‘nodeSize’. Pvalue < 0.01, enrichment > 1.75 and number of significant genes > 4 were used as thresholds to select significantly enriched GO terms. *<u>Correlation with fetal cerebellum transcriptome</u>* Transcriptome-wide correlation between bulk RNAseq data of CblOs and fetal samples was performed on fpkm-normalized expression values after selecting only protein-coding genes and filtering out lowly expressed ones (genes without at least 1 cpm in at least 3 sample). For each gene, the mean expression level at each developmental stage was calculated and correlation was computed using Spearman metrics in R. *<u>Overlap</u> <u>with NDD genes</u>* The following gene-phenotype knowledge bases were considered for the overlap analysis: (I) Development Disorder Genotype – Phenotype Database (DD2P); (II) SFARI Genes; (III) the GWAS Catalogue (EBI). SFARI and DD2P were downloaded respectively from https://gene-archive.sfari.org/tools/ and https://www.deciphergenomics.org/ddd/ddgenes. For DD2P, the overlap was tested with the complete database as well as on a subset related to the CNS, obtained by filtering the term ‘brain’ in the ‘Organ’ field. GWAS catalogue (https://www.ebi.ac.uk/gwas/) was interrogated to retrieve risk genes for 4 psychiatric disorders and two non-psychiatric conditions by searching for the indicated disease code: Attention Deficit Hyperactive disorder (EFO_0003888); Autism Spectrum Disorder (EFO_0003756); Schizophrenia (MONDO_0005090); Unipolar Depression (EFO_0003761); Diabetes Mellitus (EFO_0000400); Inflammatory Bowel Disease (EFO_0003767). Overlap significance was tested by GeneOverlap R library (version 1.32.0). Overlaps were considered significant with an odds ratio (OR) higher than 1 and P-value lower than 0.05. Results were visualized as dot plot with numbers (shared genes) shown for OR > 1, dots shown for those having also P-value < 0.05. Color-code was assigned according to OR, dot size varied according to P-value.

***CHD8*** *-* Gene count quantification at the gene level was performed using Salmon, using GRCh38 Genecode v35 as reference. A total of 13 samples were profiled: 2 wild-type and 3 mutant replicates were examined at Day 14, while at Day 21 the analysis was done in quadruplicate for both conditions. *<u>Differential expression analysis</u>* was performed comparing at each time point mutant versus wild-type organoids with DESeq2 as described above. Protein coding genes were selected for differential expression analysis. Gene filtering was applied keeping genes with at least 20 reads in at least 3 samples. Functional enrichment analysis was performed on the Molecular Function domain of the Gene Ontology by TopGO as already detailed.

### scRNAseq data pre-processing & analysis

Sample sequencing was performed on NovaSeq6000, alignment and demultiplexing of barcodes was performed using 10x Cell Ranger Multi providing gex-GRCh38-2020-A reference and Chromium_Human_Transcriptome_Probe_Set_v1.0.1_GRCh38-2020-A probe set. After demultiplexing, intra-sample doublets were detected using scDblFinder and removed. For the samples in the CHD8 dataset, the following thresholds were applied to discard low-quality droplets: minimum counts higher than 3000; minimum expressed genes higher than 2500. To discard not expressed genes, features detected in less than 1% of barcodes were discarded. For the samples in the CADPS2 dataset, mean absolute deviation (MAD)-based filter was applied to total counts (above mean+4MAD) and droplets with relatively lower read counts (below mean-1.5MAD for organoids, below mean-1.3MAD for hiPSCs), additionally, minimum number of expressing cells for a gene to be retained was set to 10 and minimum number of expressed genes for each cell was set to 200. Finally, droplets with mitochondria RNA content > 5% were excluded in both datasets. This resulted in 18116 cells and 13809 genes for the CHD8 dataset and 41639 cells and 16849 genes for the CADPS2 dataset.

All pre-processing and analyses were performed using scanpy^95^. Following normalization (target_sum=1e4 or with target_sum=2e4 for CHD8 and CADSP2 datasets, respectively) and log1p transformation, highly variable genes (HVGs) selection was carried out. For the CHD8 dataset, HGVs selection was performed using Triku^98^, while scanpy’s highly_variable_genes function was used for the CADPS2 dataset, retaining top 1500 genes, and keeping only the ones common to at least two replicates (using batch key parameter). After any dataset sub setting, HVGs selection and subsequent PCA were repeated unless differently specified. Neighborhoods graph was constructed using 65 neighbors and 20 PC for the CHD8, and 50 neighbors and 10 components for the complete dataset and 7 for the rhombic lip one (to account for differences in transcriptional heterogeneity) in the CADPS2 dataset. Diffusion maps were calculated separately for CHD8^+/E1114X^ or CHD8^+/+^ using scanpy function. For the CHD8 dataset, clusters were identified using the Leiden algorithm with a resolution of 0.6. Cluster annotation was performed through manual inspection of the top-expressed genes in each cluster, comparing them with known markers reported in the literature. To further distinguish ciliated cells with dorsal and ventral characteristics, Cluster 9 from this initial clustering was re-clustered using the Leiden algorithm at a resolution of 0.2 For the CADPS2 dataset, cell clustering was performed using Leiden algorithms with resolution parameter set to 0.8. Subsequent cell types annotation was manually curated using a combination of top expressed markers (scanpy’s rank genes function) and literature-derived markers inspection.

### Differential abundance (DA) analysis and results refinement

Differential abundance analyses were performed using Milo^52^. To compare CHD8^+/E1114X^ to CHD8^+/+^ nhoods prop was set = 0.2 and FDR threshold < 0.01. The violin plot was obtained by assigning each neighborhood to a cell type according to the most abundant one. For the rhombic lip domain in the CADPS2 dataset, PCA was corrected using harmony providing as correction covariate the sample_id to avoid highlighting differences related to technical variability among samples. After the integration, differential abundance of the corrected neighborhood’s graph was performed. Design matrix and contrast for milo were set to detect pairwise differences among genotypes (CADPS^+/+^, CADPS^Arc/Arc^ and CADPS^-/-^). To compare the CADPS^+/+^ and CADPS^Arc/Arc^, DA results were initially filtered for spatial FDR (< 0.01) absolute logFC (>1.5). Subsequently, to extract single cells belonging to enriched neighborhoods, we removed cells simultaneously belonging to discordant neighborhoods (both over and underrepresented in one condition). Finally, to label the most relevant cell types subject to differences we considered only the ones containing at least 30% of the cells (post discordance filter), belonging to enriched (or depleted) neighborhoods, resulting in Meninges_1, Late_RL and NSC_3 marked as DA cell types (**Figure 5J**).

### Differential expression analysis

Differentially expressed genes were identified by comparing CHD8^+/E1114X^ or CHD8^+/+^ using the Wilcoxon test implementation in scanpy on cells of EN/iGN clusters. Genes were then ranked by fold-change and functional enrichment analysis was tested by gseapy on Reactome gene sets. Transcription factor activity for Catenin B was calculated by Multivariate Linear Model in decoupleR^99^ using Collect3 as database.

### Functional analysis of enriched domains

The three identified DA cell types were subjected to functional analyses to extract transcriptional programs that sharply characterize the differences among CADPS2^+/+^ and CADPS2^Arc/Arc^. Meninges_1, Late_RL and NSCs_3 domains were grouped by cell type (**Figure 5I**) and enrichment status (**Figure 5K**) obtaining the following contrasts for DEA: Meninges_1: CADPS2^Arc/Arc^ -enriched *vs.* Neutral, Late_RL: CADPS2^+/+^ - enriched *vs*. Neutral. For NSC_3: CADPS2^Arc/Arc^ -enriched *vs.* CADPS2^+/+^-enriched, we took advantage of symmetrically opposite enriched cells neighborhoods within the same cell type. The defined cell groups were subsampled, when needed, to ensure balance among them, and pseudobulk approach was adopted to random aggregate cells into five pseudo replicates *per* group. edgeR^100^ GLM was applied to implement differential expression analysis. Functional analysis was subsequently performed using gProfiler^101^ python implementation providing the deriving DE genes for each contrast (FDR < 0.01, absolute logFC > 1.5) and as background the genes expressed in at least *n*-1 pseudoreplicates for each considered contrast. Terms were additionally filtered for minimum overlap of 10 genes. For CADPS2 dataset, hiPSCs were firstly downsampled, when needed, to bring the four compared groups (CADPS2^Arc/Arc^ and CADPS2^+/+^ at 3% or 21% O_2_) to the same number of cells by cell cycle phase to avoid DEA results being driven by any imbalance. Also, in this case pseudobulk approach was adopted prior to differential expression, targeting five pseudoreplicates per group. DEA was run via edgeR quasi-likelihood negative binomial generalized log-linear model (glmQLFit) after operating normalization and dispersion estimation using calcNormFactors, and estimateGLMRobustDispersion respectively. Deriving genes were considered differentially expressed if FDR < 0.01 and absolute logFC > 1.5.

### ChIP-seq analysis

We performed ChIPseq reads alignment for HIF1a on the human hg38 genome using Bowtie 1.0 (-v 2 - m 1). Peak calling was performed with MACS2, using narrow parameters (-q 0.1). To perform ChIPseq quantitative analysis, including PCA, we selected the peaks of each mark identified in at least two samples (independently of their genotype) and measured read-counts *per* peak in each sample with DeepTools multiBamSummary. We performed qualitative analyses by grouping: i) peaks found in all the samples from a specific condition and present in at most n=1 sample of the other condition (e.g. - CADPS2^Arc/Arc^ *vs* CADPS2^+/+^ or CADPS2^+/+ HiO2^ *vs* CADPS2^+/+ LowO2^), with ii) peaks found in at least n=1 samples of a certain genotype and not present in the other. Both groupings were performed with BedTools (version 2.26), MultiIntersect function. Reference peaks were defined as present in both control samples. We identified “lost peaks” as regions preferentially found in one condition (e.g., regions in at least n=1 controls and none of mutant line, plus regions found in all controls and at most n=1 mutant). Peaks were considered “gained” following the inverse logic.

## STATISTICS

RStudio was used to perform statistical analysis of bioinformatics experiments. All other statistical tests were performed with Prism10. In each figure legend, results are indicated as mean ± SEM along with sample size, number of experimental replicates as well as the statistical test applied and the resulting p-values. Statistical significance was defined as a p-value < 0.05 with ∗ < 0.05, ∗∗ < 0.01, and ∗∗∗ < 0.001. Imaging analyses were performed in blind.

## Notes

### Competing Interest Statement

The authors have declared no competing interest.

## REFERENCES

1. Rakic, P., and Sidman, R.L. (1970). Histogenesis of cortical layers in human cerebellum, particularly the lamina dissecans. Journal of Comparative Neurology 139, 473–500. 10.1002/CNE.901390407.

2. Haldipur, P., Millen, K.J., and Aldinger, K.A. (2022). Human Cerebellar Development and Transcriptomics: Implications for Neurodevelopmental Disorders. Annu Rev Neurosci 45, 515–531. 10.1146/ANNUREV-NEURO-111020-091953/CITE/REFWORKS.

3. 3. van Essen, M.J., Nayler, S., Becker, E.B.E., and Jacob, J. (2020). Deconstructing cerebellar development cell by cell. PLoS Genet 16, e1008630. 10.1371/JOURNAL.PGEN.1008630.

4. Robinson, K.J., Watchon, M., and Laird, A.S. (2020). Aberrant Cerebellar Circuitry in the Spinocerebellar Ataxias. Front Neurosci 14, 559743. 10.3389/FNINS.2020.00707/PDF.

5. Hendrikse, L.D., Haldipur, P., Saulnier, O., Millman, J., Sjoboen, A.H., Erickson, A.W., Ong, W., Gordon, V., Coudière-Morrison, L., Mercier, A.L., et al. (2022). Failure of human rhombic lip differentiation underlies medulloblastoma formation. Nature 2022 609:7929 *609*, 1021–1028. 10.1038/s41586-022-05215-w.

6. Aldinger, K.A., Thomson, Z., Phelps, I.G., Haldipur, P., Deng, M., Timms, A.E., Hirano, M., Santpere, G., Roco, C., Rosenberg, A.B., et al. (2021). Spatial and cell type transcriptional landscape of human cerebellar development. Nature Neuroscience 2021 24:8 *24*, 1163–1175. 10.1038/s41593-021-00872-y.

7. Becker, E.B.E., and Stoodley, C.J. (2013). Autism Spectrum Disorder and the Cerebellum. Int Rev Neurobiol 113, 1–34. 10.1016/B978-0-12-418700-9.00001-0.

8. Wang, S.S.H., Kloth, A.D., and Badura, A. (2014). The Cerebellum, Sensitive Periods, and Autism. Neuron 83, 518–532. 10.1016/J.NEURON.2014.07.016/ASSET/386BF1FD-E929-436D-99E7-F9F90D238341/MAIN.ASSETS/GR4.JPG.

9. Krishnan, A., Zhang, R., Yao, V., Theesfeld, C.L., Wong, A.K., Tadych, A., Volfovsky, N., Packer, A., Lash, A., and Troyanskaya, O.G. (2016). Genome-wide prediction and functional characterization of the genetic basis of autism spectrum disorder. Nature Neuroscience 2016 19:11 *19*, 1454–1462. 10.1038/nn.4353.

10. Stoodley, C.J. (2016). The Cerebellum and Neurodevelopmental Disorders. Cerebellum 15, 34–37. 10.1007/S12311-015-0715-3/METRICS.

11. Romer, A.L., Knodt, A.R., Houts, R., Brigidi, B.D., Moffitt, T.E., Caspi, A., and Hariri, A.R. (2017). Structural alterations within cerebellar circuitry are associated with general liability for common mental disorders. Molecular Psychiatry 2018 23:4 *23*, 1084–1090. 10.1038/mp.2017.57.

12. d’Oleire Uquillas, F., Sefik, E., Li, B., Trotter, M.A., Steele, K.A., Seidlitz, J., Gesue, R., Latif, M., Fasulo, T., Zhang, V., et al. (2024). Multimodal evidence for cerebellar influence on cortical development in autism: structural growth amidst functional disruption. Molecular Psychiatry 2024, 1–15. 10.1038/s41380-024-02769-1.

13. Zhang, R., Zhou, X., Yuan, D., Lu, Q., Chen, X., and Zhang, Y. (2025). Associations between cerebellum and major psychiatric disorders: a bidirectional Mendelian randomization study. European Archives of Psychiatry and Clinical Neuroscience 2025, 1–11. 10.1007/S00406-025-01971-8.

14. Hublin, J.J., Neubauer, S., and Gunz, P. (2015). Brain ontogeny and life history in Pleistocene hominins. Philosophical Transactions of the Royal Society B: Biological Sciences 370. 10.1098/RSTB.2014.0062.

15. Barton, R.A., and Venditti, C. (2014). Rapid evolution of the cerebellum in humans and other great apes. Current Biology 24, 2440–2444. 10.1016/j.cub.2014.08.056.

16. Buisan, R., Moriano, J., Andirkó, A., and Boeckx, C. (2022). A Brain Region-Specific Expression Profile for Genes Within Large Introgression Deserts and Under Positive Selection in Homo sapiens. Front Cell Dev Biol 10, 824740. 10.3389/FCELL.2022.824740/BIBTEX.

17. Rickelton, K., Ely, J.J., Hopkins, W.D., Hof, P.R., Sherwood, C.C., Bauernfeind, A.L., and Babbitt, C.C. (2025). Transcriptomic changes across subregions of the primate cerebellum support the evolution of uniquely human behaviors. bioRxiv, 2025.03.03.641249. 10.1101/2025.03.03.641249.

18. Rickelton, K., Zintel, T.M., Pizzollo, J., Miller, E., Ely, J.J., Raghanti, M.A., Hopkins, W.D., Hof, P.R., Sherwood, C.C., Bauernfeind, A.L., et al. (2024). Tempo and mode of gene expression evolution in the brain across primates. Elife 13. 10.7554/ELIFE.70276.

19. Wang, B., LeBel, A., and D’Mello, A.M. (2025). Ignoring the cerebellum is hindering progress in neuroscience. Trends Cogn Sci. 10.1016/J.TICS.2025.01.004.

20. Pașca, S.P., Arlotta, P., Bateup, H.S., Camp, J.G., Cappello, S., Gage, F.H., Knoblich, J.A., Kriegstein, A.R., Lancaster, M.A., Ming, G.L., et al. (2022). A nomenclature consensus for nervous system organoids and assembloids. Nature 2022 609:7929 *609*, 907–910. 10.1038/s41586-022-05219-6.

21. Cheroni, C., Trattaro, S., Caporale, N., López-Tobón, A., Tenderini, E., Sebastiani, S., Troglio, F., Gabriele, M., Bressan, R.B., Pollard, S.M., et al. (2022). Benchmarking brain organoid recapitulation of fetal corticogenesis. Translational Psychiatry 2022 12:1 *12*, 1–16. 10.1038/s41398-022-02279-0.

22. Atamian, A., Birtele, M., Hosseini, N., Nguyen, T., Seth, A., Del Dosso, A., Paul, S., Tedeschi, N., Taylor, R., Coba, M.P., et al. (2024). Human cerebellar organoids with functional Purkinje cells. Cell Stem Cell 31, 39–51.e6. 10.1016/j.stem.2023.11.013.

23. Amin, N.D., Kelley, K.W., Kaganovsky, K., Onesto, M., Hao, J., Miura, Y., McQueen, J.P., Reis, N., Narazaki, G., Li, T., et al. (2024). Generating human neural diversity with a multiplexed morphogen screen in organoids. Cell Stem Cell 31, 1831–1846.e9. 10.1016/J.STEM.2024.10.016.

24. Muguruma, K., Nishiyama, A., Kawakami, H., Hashimoto, K., and Sasai, Y. (2015). Self-Organization of Polarized Cerebellar Tissue in 3D Culture of Human Pluripotent Stem Cells. Cell Rep 10, 537–550. 10.1016/J.CELREP.2014.12.051.

25. Nayler, S., Agarwal, D., Curion, F., Bowden, R., and Becker, E.B.E. (2021). High-resolution transcriptional landscape of xeno-free human induced pluripotent stem cell-derived cerebellar organoids. 11, 1–17.

26. Silva, T.P., Sousa-Luís, R., Fernandes, T.G., Bekman, E.P., Rodrigues, C.A.V., Vaz, S.H., Moreira, L.M., Hashimura, Y., Jung, S., Lee, B., et al. (2021). Transcriptome profiling of human pluripotent stem cell-derived cerebellar organoids reveals faster commitment under dynamic conditions. Biotechnol Bioeng 118, 2781–2803. 10.1002/BIT.27797.

27. Ballabio, C., Anderle, M., Gianesello, M., Lago, C., Miele, E., Cardano, M., Aiello, G., Piazza, S., Caron, D., Gianno, F., et al. (2020). Modeling medulloblastoma in vivo and with human cerebellar organoids. Nature Communications 2020 11:1 *11*, 1–18. 10.1038/s41467-019-13989-3.

28. 28. van Essen, M.J., Apsley, E.J., Riepsaame, J., Xu, R., Northcott, P.A., Cowley, S.A., Jacob, J., and Becker, E.B.E. (2024). PTCH1-mutant human cerebellar organoids exhibit altered neural development and recapitulate early medulloblastoma tumorigenesis. DMM Disease Models and Mechanisms 17. 10.1242/DMM.050323/344006.

29. Kamei, T., Tamada, A., Kimura, T., Kakizuka, A., Asai, A., and Muguruma, K. (2023). Survival and process outgrowth of human iPSC-derived cells expressing Purkinje cell markers in a mouse model for spinocerebellar degenerative disease. Exp Neurol 369, 114511. 10.1016/J.EXPNEUROL.2023.114511.

30. Pattabiraman, K., Muchnik, S.K., and Sestan, N. (2020). The evolution of the human brain and disease susceptibility. Curr Opin Genet Dev 65, 91–97. 10.1016/J.GDE.2020.05.004.

31. O ’ Roak, B.J., Vives, L., Girirajan, S., Karakoc, E., Krumm, N., Coe, B.P., Levy, R., Ko, A., Lee, C., Smith, J.D., et al. (2012). Sporadic autism exomes reveal a highly interconnected protein network of de novo mutations. Nature 2012 485:7397 *485*, 246–250. 10.1038/nature10989.

32. O’Roak, B.J., Vives, L., Fu, W., Egertson, J.D., Stanaway, I.B., Phelps, I.G., Carvill, G., Kumar, A., Lee, C., Ankenman, K., et al. (2012). Multiplex targeted sequencing identifies recurrently mutated genes in autism spectrum disorders. Science (1979) *338*, 1619–1622. 10.1126/SCIENCE.1227764/SUPPL_FILE/OROAK.SM.PDF.

33. 33. Bernier, R., Golzio, C., Xiong, B., Stessman, H.A., Coe, B.P., Penn, O., Witherspoon, K., Gerdts, J., Baker, C., Vulto-Van Silfhout, A.T., et al. (2014). Disruptive CHD8 Mutations Define a Subtype of Autism Early in Development. Cell 158, 263–276. 10.1016/J.CELL.2014.06.017.

34. Bonora, E., Graziano, C., Minopoli, F., Bacchelli, E., Magini, P., Diquigiovanni, C., Lomartire, S., Bianco, F., Vargiolu, M., Parchi, P., et al. (2014). Maternally inherited genetic variants of CADPS2 are present in Autism Spectrum Disorders and Intellectual Disability patients. EMBO Mol Med 6, 795–809. 10.1002/EMMM.201303235/SUPPL_FILE/EMMM201303235.R EVIEWER_COMMENTS.PDF.

35. Rovira, P., Demontis, D., Sánchez-Mora, C., Zayats, T., Klein, M., Mota, N.R., Weber, H., Garcia-Martínez, I., Pagerols, M., Vilar-Ribó, L., et al. (2020). Shared genetic background between children and adults with attention deficit/hyperactivity disorder. Neuropsychopharmacology 2020 45:10 *45*, 1617–1626. 10.1038/s41386-020-0664-5.

36. Kullmann, J.A., Trivedi, N., Howell, D., Laumonnerie, C., Nguyen, V., Banerjee, S.S., Stabley, D.R., Shirinifard, A., Rowitch, D.H., and Solecki, D.J. (2020). Oxygen Tension and the VHL-Hif1α Pathway Determine Onset of Neuronal Polarization and Cerebellar Germinal Zone Exit. Neuron 106, 607–623.e5.

37. Sollis, E., Mosaku, A., Abid, A., Buniello, A., Cerezo, M., Gil, L., Groza, T., Güneş, O., Hall, P., Hayhurst, J., et al. (2023). The NHGRI-EBI GWAS Catalog: knowledgebase and deposition resource. Nucleic Acids Res 51, D977–D985. 10.1093/NAR/GKAC1010.

38. Villa, C.E., Cheroni, C., Dotter, C.P., López-Tóbon, A., Oliveira, B., Sacco, R., Yahya, A.Ç., Morandell, J., Gabriele, M., Tavakoli, M.R., et al. (2022). CHD8 haploinsufficiency links autism to transient alterations in excitatory and inhibitory trajectories. Cell Rep 39, 110615. 10.1016/J.CELREP.2022.110615.

39. Kawamura, A., Katayama, Y., Kakegawa, W., Ino, D., Nishiyama, M., Yuzaki, M., and Nakayama, K.I. (2021). The autism-associated protein CHD8 is required for cerebellar development and motor function. Cell Rep 35. 10.1016/J.CELREP.2021.108932.

40. Rodríguez-Paredes, M., Ceballos-Chávez, M., Esteller, M., García-Domínguez, M., and Reyes, J.C. (2009). The chromatin remodeling factor CHD8 interacts with elongating RNA polymerase II and controls expression of the cyclin E2 gene. Nucleic Acids Res 37, 2449–2460. 10.1093/NAR/GKP101.

41. Sood, S., Webera, C.M., Hodges, H.C., Krokhotin, A., Shalizi, A., and Crabtree, G.R. (2020). CHD8 dosage regulates transcription in pluripotency and early murine neural differentiation. Proc Natl Acad Sci U S A 117, 22331. 10.1073/PNAS.1921963117/SUPPL_FILE/PNAS.1921963117.S APP.PDF.

42. Nishiyama, M., Skoultchi, A.I., and Nakayama, K.I. (2012). Histone H1 Recruitment by CHD8 Is Essential for Suppression of the Wnt–β-Catenin Signaling Pathway. Mol Cell Biol 32, 501–512. 10.1128/MCB.06409-11.

43. Nishiyama, M., Nakayama, K., Tsunematsu, R., Tsukiyama, T., Kikuchi, A., and Nakayama, K.I. (2004). Early Embryonic Death in Mice Lacking the β-Catenin-Binding Protein Duplin. Mol Cell Biol 24, 8386–8394. 10.1128/MCB.24.19.8386-8394.2004.

44. Durak, O., Gao, F., Kaeser-Woo, Y.J., Rueda, R., Martorell, A.J., Nott, A., Liu, C.Y., Watson, L.A., and Tsai, L.H. (2016). Chd8 mediates cortical neurogenesis via transcriptional regulation of cell cycle and Wnt signaling. Nature Neuroscience 2016 19:11 *19*, 1477–1488. 10.1038/nn.4400.

45. Wang, P., Mokhtari, R., Pedrosa, E., Kirschenbaum, M., Bayrak, C., Zheng, D., and Lachman, H.M. (2017). CRISPR/Cas9-mediated heterozygous knockout of the autism gene CHD8 and characterization of its transcriptional networks in cerebral organoids derived from iPS cells. Molecular Autism 2017 8:1 *8*, 1–17. 10.1186/S13229-017-0124-1.

46. Schüller, U., and Rowitch, D.H. (2007). β-catenin function is required for cerebellar morphogenesis. Brain Res 1140, 161–169. 10.1016/J.BRAINRES.2006.05.105.

47. Selvadurai, H.J., and Mason, J.O. (2011). Wnt/β-catenin Signalling Is Active in a Highly Dynamic Pattern during Development of the Mouse Cerebellum. 6, e23012. 10.1371/JOURNAL.PONE.0023012.

48. Lasser, M., Sun, N., Xu, Y., Wang, S., Drake, S., Law, K., Gonzalez, S., Wang, B., Drury, V., Castillo, O., et al. (2023). Pleiotropy of autism-associated chromatin regulators. Development (Cambridge) 150. 10.1242/DEV.201515/319900/AM/PLEIOTROPY-OF-AUTISM-ASSOCIATED-CHROMATIN.

49. Igreja, L., Menezes, C., Pinto, P.S., Freixo, J.P., and Chorão, R. (2023). Lissencephaly With Cerebellar Hypoplasia Due To a New RELN Mutation. Pediatr Neurol 149, 137–140. 10.1016/J.PEDIATRNEUROL.2023.09.012.

50. Valence, S., Garel, C., Barth, M., Toutain, A., Paris, C., Amsallem, D., Barthez, M.A., Mayer, M., Rodriguez, D., and Burglen, L. (2016). RELN and VLDLR mutations underlie two distinguishable clinico-radiological phenotypes. Clin Genet 90, 545–549. 10.1111/CGE.12779.

51. Coutelier, M., Burglen, L., Mundwiller, E., Abada-Bendib, M., Rodriguez, D., Chantot-Bastaraud, S., Rougeot, C., Cournelle, M.A., Milh, M., Toutain, A., et al. (2015). GRID2 mutations span from congenital to mild adult-onset cerebellar ataxia. Neurology 84, 1751–1759. 10.1212/WNL.0000000000001524/SUPPL_FILE/FIGURE_E-1.DOCX.

52. Dann, E., Henderson, N.C., Teichmann, S.A., Morgan, M.D., and Marioni, J.C. (2021). Differential abundance testing on single-cell data using k-nearest neighbor graphs. Nature Biotechnology 2021 40:2 *40*, 245–253. 10.1038/s41587-021-01033-z.

53. Jho, E., Zhang, T., Domon, C., Joo, C.-K., Freund, J.-N., and Costantini, F. (2002). Wnt/β-Catenin/Tcf Signaling Induces the Transcription of Axin2, a Negative Regulator of the Signaling Pathway. Mol Cell Biol 22, 1172–1183. 10.1128/MCB.22.4.1172-1183.2002.

54. Glinka, A., Wu, W., Delius, H., Monaghan, A.P., Blumenstock, C., and Niehrs, C. (1998). Dickkopf-1 is a member of a new family of secreted proteins and functions in head induction. Nature 1998 391:6665 *391*, 357–362. 10.1038/34848.

55. Huggins, I.J., Bos, T., Gaylord, O., Jessen, C., Lonquich, B., Puranen, A., Richter, J., Rossdam, C., Brafman, D., Gaasterland, T., et al. (2017). The WNT target SP5 negatively regulates WNT transcriptional programs in human pluripotent stem cells. Nature Communications 2017 8:1 *8*, 1–14. 10.1038/s41467-017-01203-1.

56. Pei, Y., Brun, S.N., Markant, S.L., Lento, W., Gibson, P., Taketo, M.M., Giovannini, M., Gilbertson, R.J., and Wechsler-Reya, R.J. (2012). WNT signaling increases proliferation and impairs differentiation of stem cells in the developing cerebellum. Development 139, 1724–1733. 10.1242/DEV.050104.

57. Rubenstein, J.L.R., and Merzenich, M.M. (2003). Model of autism: increased ratio of excitation/inhibition in key neural systems. Genes Brain Behav 2, 255–267. 10.1034/J.1601-183X.2003.00037.X.

58. Cellot, G., and Cherubini, E. (2014). GABAergic signaling as therapeutic target for autism spectrum disorders. Front Pediatr 2, 102071. 10.3389/FPED.2014.00070/BIBTEX.

59. Kuhlwilm, M., and Boeckx, C. (2019). A catalog of single nucleotide changes distinguishing modern humans from archaic hominins. Scientific Reports 2019 9:1 *9*, 1–14. 10.1038/s41598-019-44877-x.

60. Moriano, J., Leonardi, O., Vitriolo, A., Testa, G., and Boeckx, C. (2024). A multi-layered integrative analysis reveals a cholesterol metabolic program in outer radial glia with implications for human brain evolution. Development (Cambridge) 151. 10.1242/DEV.202390/361499/AM/A-MULTI-LAYERED-INTEGRATIVE-ANALYSIS-REVEALS-A.

61. Peyregne, S., Boyle, M.J., Dannemann, M., and Prufer, K. (2017). Detecting ancient positive selection in humans using extended lineage sorting. Genome Res 27, 1563–1572. 10.1101/GR.219493.116/-/DC1.

62. Vernot, B., Tucci, S., Kelso, J., Schraiber, J.G., Wolf, A.B., Gittelman, R.M., Dannemann, M., Grote, S., McCoy, R.C., Norton, H., et al. (2016). Excavating Neandertal and Denisovan DNA from the genomes of Melanesian individuals. Science (1979) 352, 235–239. 10.1126/SCIENCE.AAD9416/SUPPL_FILE/VERNOT-SM.PDF.

63. McLean, C.Y., Bristor, D., Hiller, M., Clarke, S.L., Schaar, B.T., Lowe, C.B., Wenger, A.M., and Bejerano, G. (2010). GREAT improves functional interpretation of cis-regulatory regions. Nature Biotechnology 2010 28:5 *28*, 495–501. 10.1038/nbt.1630.

64. Cao, Y., Wang, X., and Peng, G. (2020). SCSA: A cell type annotation tool for single-cell RNA-seq data. Front Genet 11, 524690. 10.3389/FGENE.2020.00490/BIBTEX.

65. 65. Yu, C., Liu, Y., Ma, T., Liu, K., Xu, S., Zhang, Y., Liu, H., La Russa, M., Xie, M., Ding, S., et al. (2015). Small molecules enhance crispr genome editing in pluripotent stem cells. Cell Stem Cell 16, 142–147. 10.1016/j.stem.2015.01.003.

66. Jacquet, B. V., Salinas-Mondragon, R., Liang, H., Therit, B., Buie, J.D., Dykstra, M., Campbell, K., Ostrowski, L.E., Brody, S.L., and Ghashghaei, H.T. (2009). FoxJ1-dependent gene expression is required for differentiation of radial glia into ependymal cells and a subset of astrocytes in the postnatal brain. Development 136, 4021–4031. 10.1242/DEV.041129.

67. Chang, C.H., Zanini, M., Shirvani, H., Cheng, J.S., Yu, H., Feng, C.H., Mercier, A.L., Hung, S.Y., Forget, A., Wang, C.H., et al. (2019). Atoh1 Controls Primary Cilia Formation to Allow for SHH-Triggered Granule Neuron Progenitor Proliferation. Dev Cell 48, 184–199.e5. 10.1016/J.DEVCEL.2018.12.017/ATTACHMENT/E046ECE6-DF0B-407C-915D-E27F6DA9C98D/MMC2.PDF.

68. Hellwig, M., Lauffer, M.C., Bockmayr, M., Spohn, M., Merk, D.J., Harrison, L., Ahlfeld, J., Kitowski, A., Neumann, J.E., Ohli, J., et al. (2019). TCF4 (E2-2) harbors tumor suppressive functions in SHH medulloblastoma. Acta Neuropathol 137, 657–673. 10.1007/S00401-019-01982-5/METRICS.

69. Chodelkova, O., Masek, J., Korinek, V., Kozmik, Z., and Machon, O. (2018). Tcf7L2 is essential for neurogenesis in the developing mouse neocortex. Neural Dev 13, 1–10. 10.1186/S13064-018-0107-8/FIGURES/5.

70. Hryniewiecka, K., Lipiec, M., Majkowska, M., Baggio, S., Kublik, E., and Wiśniewska, M. (2023). THE ROLE OF TCF7L2 TRANSCRIPTION FACTOR IN THE FUCTION OF THALAMO-CORTICAL CIRCUITS AND IN PATHOGENESIS OF AUTISM SPECTRUM DISORDER. IBRO Neurosci Rep 15, S102. 10.1016/J.IBNEUR.2023.08.085.

71. Imitola, J., Hollingsworth, E.W., Watanabe, F., Olah, M., Elyaman, W., Starossom, S., Kivis&#x00E4;kk, P., and Khoury, S.J. (2023). Stat1 is an inducible transcriptional repressor of neural stem cells self-renewal program during neuroinflammation. Front Cell Neurosci 17, 1156802. 10.3389/FNCEL.2023.1156802/BIBTEX.

72. Ferluga, S., Baiz, D., Hilton, D.A., Adams, C.L., Ercolano, E., Dunn, J., Bassiri, K., Kurian, K.M., and Hanemann, C.O. (2020). Constitutive activation of the EGFR–STAT1 axis increases proliferation of meningioma tumor cells. Neurooncol Adv 2, 1–12. 10.1093/NOAJNL/VDAA008.

73. Krycer, J.R., and Nayler, S.P. (2022). A Survey of the Metabolic Landscape of the Developing Cerebellum at Single-Cell Resolution. Cerebellum 21, 838– 850. 10.1007/S12311-022-01415-2/METRICS.

74. Haldipur, P., Aldinger, K.A., Bernardo, S., Deng, M., Timms, A.E., Overman, L.M., Winter, C., Lisgo, S.N., Razavi, F., Silvestri, E., et al. (2019). Spatiotemporal expansion of primary progenitor zones in the developing human cerebellum. Science (1979) 366, 454–460. 10.1126/SCIENCE.AAX7526/SUPPL_FILE/AAX7526_HALDIPU R_SM_REV.PDF.

75. Hartmann, D., Ziegenhagen, M.W., and Sievers, J. (1998). Meningeal cells stimulate neuronal migration and the formation of radial glial fascicles from the cerebellar external granular layer. Neurosci Lett 244, 129–132. 10.1016/S0304-3940(98)00154-2.

76. Reiss, K., Mentlein, R., Sievers, J., and Hartmann, D. (2002). Stromal cell-derived factor 1 is secreted by meningeal cells and acts as chemotactic factor on neuronal stem cells of the cerebellar external granular layer. Neuroscience 115, 295–305. 10.1016/S0306-4522(02)00307-X.

77. Zhu, Y., Yu, T., and Rao, Y. (2004). Temporal regulation of cerebellar EGL migration through a switch in cellular responsiveness to the meninges. Dev Biol 267, 153–164. 10.1016/J.YDBIO.2003.10.037.

78. Haldipur, P., Gillies, G.S., Janson, O.K., Chizhikov, V. V., Mithal, D.S., Miller, R.J., and Millen, K.J. (2014). Foxc1 dependent mesenchymal signalling drives embryonic cerebellar growth. Elife 3. 10.7554/ELIFE.03962.

79. Borrell, V., and Marín, O. (2006). Meninges control tangential migration of hem-derived Cajal-Retzius cells via CXCL12/CXCR4 signaling. Nature Neuroscience 2006 9:10 *9*, 1284–1293. 10.1038/nn1764.

80. Siegenthaler, J.A., and Pleasure, S.J. (2011). We have got you ‘covered’: how the meninges control brain development. Curr Opin Genet Dev 21, 249–255. 10.1016/J.GDE.2010.12.005.

81. Como, C.N., Kim, S., and Siegenthaler, J. (2023). Stuck on you: Meninges cellular crosstalk in development. Curr Opin Neurobiol 79, 102676. 10.1016/J.CONB.2023.102676.

82. Gordon, A., Yoon, S.-J., Bicks, L.K., Martin, J.M., Pintacuda, G., Arteaga, S., Wamsley, B., Guo, Q., Elahi, L., Dolmetsch, R.E., et al. (2024). Developmental convergence and divergence in human stem cell models of autism spectrum disorder. bioRxiv, 2024.04.01.587492. 10.1101/2024.04.01.587492.

83. Chen, X., Chen, T., Dong, C., Chen, H., Dong, X., Yang, L., Hu, L., Wang, H., Wu, B., Yao, Y., et al. (2022). Deletion of CHD8 in cerebellar granule neuron progenitors leads to severe cerebellar hypoplasia, ataxia, and psychiatric behavior in mice. 49, 859–869. 10.1016/J.JGG.2022.02.011.

84. Li, B., Zhao, H., Tu, Z., Yang, W., Han, R., Wang, L., Luo, X., Pan, M., Chen, X., Zhang, J., et al. (2023). CHD8 mutations increase gliogenesis to enlarge brain size in the nonhuman primate. 9, 1–23.

85. Kawamura, A., Katayama, Y., Nishiyama, M., Shoji, H., Tokuoka, K., Ueta, Y., Miyata, M., Isa, T., Miyakawa, T., Hayashi-Takagi, A., et al. (2020). Oligodendrocyte dysfunction due to Chd8 mutation gives rise to behavioral deficits in mice. Hum Mol Genet 29, 1274–1291. 10.1093/HMG/DDAA036.

86. Jin, X., Simmons, S.K., Guo, A., Shetty, A.S., Ko, M., Nguyen, L., Jokhi, V., Robinson, E., Oyler, P., Curry, N., et al. (2020). In vivo Perturb-Seq reveals neuronal and glial abnormalities associated with autism risk genes. Science (1979) 370. 10.1126/SCIENCE.AAZ6063/SUPPL_FILE/AAZ6063_JIN_TABLE-S9.XLSX.

87. Molz, B., Alberro, M.L., Eising, E., Schijven, D., Alagöz, G., Francks, C., and Fisher, S.E. (2024). Evaluating the effects of archaic protein-altering variants in living human adults. bioRxiv, 2024.07.05.602242. 10.1101/2024.07.05.602242.

88. Takagi, K., and Kondo, O. (2025). Postnatal interaction of size and shape in the human endocranium and brain structures. bioRxiv, 2025.01.20.633848. 10.1101/2025.01.20.633848.

89. Funato, N., Heliövaara, A., and Boeckx, C. (2024). A regulatory variant impacting TBX1 expression contributes to basicranial morphology in Homo sapiens. The American Journal of Human Genetics 111, 939–953. 10.1016/J.AJHG.2024.03.012.

90. Iwata, R., and Vanderhaeghen, P. (2024). Metabolic mechanisms of species-specific developmental tempo. Dev Cell 59, 1628–1639. 10.1016/J.DEVCEL.2024.05.027.

91. Kshetri, R., Beavers, J.O., Hyde, R., Ewa, R., Schwertman, A., Porcayo, S., and Richardson, B.D. (2024). Behavioral decline in Shank3Δex4–22 mice during early adulthood parallels cerebellar granule cell glutamatergic synaptic changes. Molecular Autism 2024 15:1 *15*, 1–21. 10.1186/S13229-024-00628-Y.

92. Zeberg, H., Jakobsson, M., and Pääbo, S. (2024). The genetic changes that shaped Neandertals, Denisovans, and modern humans. Cell 187, 1047–1058. 10.1016/J.CELL.2023.12.029/ASSET/6466FCBC-2896-43B2-92B9-6BE976CAD007/MAIN.ASSETS/GR4.JPG.

93. Caporale, N., Leonardi, O., Villa, C.E., Vitriolo, A., Boeckx, C., and Testa, G. (2025). Tile by tile: capturing the evolutionary mosaic of human conditions. Curr Opin Genet Dev 90, 102297. 10.1016/J.GDE.2024.102297.

94. Maccione, A., Gandolfo, M., Massobrio, P., Novellino, A., Martinoia, S., and Chiappalone, M. (2009). A novel algorithm for precise identification of spikes in extracellularly recorded neuronal signals. J Neurosci Methods 177, 241–249. 10.1016/J.JNEUMETH.2008.09.026.

95. Wolf, F.A., Angerer, P., and Theis, F.J. (2018). SCANPY: Large-scale single-cell gene expression data analysis. Genome Biol 19, 1–5. 10.1186/S13059-017-1382-0/FIGURES/1.

96. Pereira, M.F., Finazzi, V., Rizzuti, L., Aprile, D., Aiello, V., Mollica, L., Riva, M., Soriani, C., Dossena, F., Shyti, R., et al. (2025). YY1 mutations disrupt corticogenesis through a cell type specific rewiring of cell-autonomous and non-cell-autonomous transcriptional programs. Molecular Psychiatry 2025, 1–17. 10.1038/s41380-025-02929-x.

97. Love, M.I., Huber, W., and Anders, S. (2014). Moderated estimation of fold change and dispersion for RNA-seq data with DESeq2. Genome Biol 15, 1–21. 10.1186/S13059-014-0550-8/FIGURES/9.

98. M Ascensión, A., Ibáñez-SolCrossed D Sign©, O., Inza, I., Izeta, A., and Araúzo-Bravo, M.J. (2022). Triku: a feature selection method based on nearest neighbors for single-cell data. Gigascience 11, 1–16. 10.1093/GIGASCIENCE/GIAC017.

99. Badia-I-Mompel, P., Vélez Santiago, J., Braunger, J., Geiss, C., Dimitrov, D., Müller-Dott, S., Taus, P., Dugourd, A., Holland, C.H., Ramirez Flores, R.O., et al. (2022). decoupleR: ensemble of computational methods to infer biological activities from omics data. Bioinformatics Advances 2. 10.1093/BIOADV/VBAC016.

100. Chen, Y., Chen, L., Lun, A.T.L., Baldoni, P.L., and Smyth, G.K. (2025). edgeR v4: powerful differential analysis of sequencing data with expanded functionality and improved support for small counts and larger datasets. Nucleic Acids Res 53, 13–14. 10.1093/NAR/GKAF018.

101. Kolberg, L., Raudvere, U., Kuzmin, I., Adler, P., Vilo, J., and Peterson, H. (2023). g:Profiler—interoperable web service for functional enrichment analysis and gene identifier mapping (2023 update). Nucleic Acids Res 51, W207–W212. 10.1093/NAR/GKAD347.

