## Supplementary Figure 1 for "Benchmarking cerebellar organoids to model autism spectrum disorder and human brain evolution"

### Figure S1

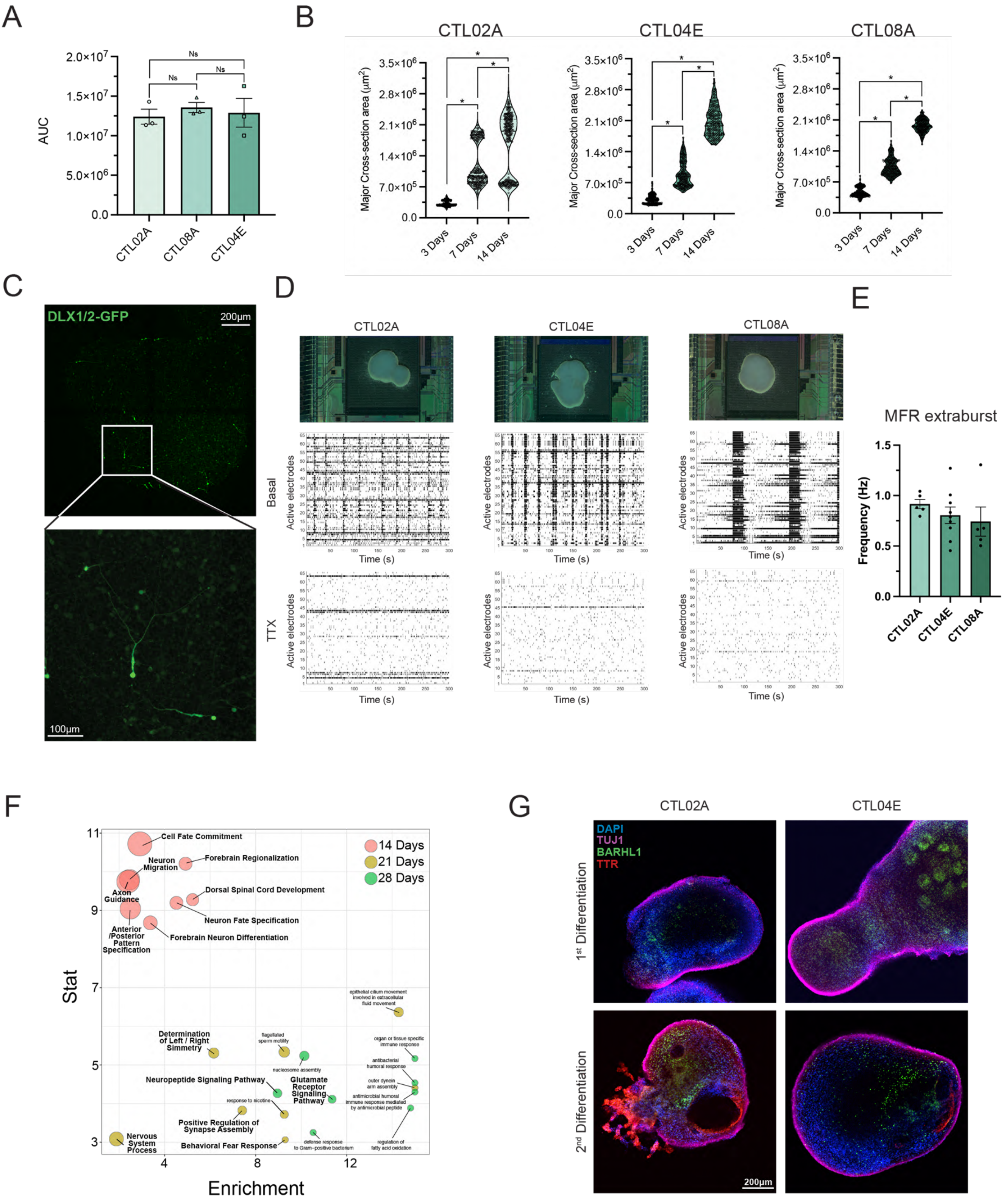
A
AUC
Ns
Ns
Ns
CTL02A
CTL08A
CTL04E
B
CTL02A
CTL04E
CTL08A
Major Cross-section area (μm²)
3 Days
7 Days
14 Days
C
DLX1/2-GFP
200μm
100μm
D
CTL02A
CTL04E
CTL08A
Basal
TTX
Active electrodes
Time (s)
E
MFR extraburst
Frequency (Hz)
CTL02A
CTL04E
CTL08A
F
Stat
Enrichment
14 Days
21 Days
28 Days
Cell Fate Commitment
Forebrain Regionalization
Neuron Migration
Axon Guidance
Dorsal Spinal Cord Development
Neuron Fate Specification
Anterior /Posterior Pattern Specification
Forebrain Neuron Differentiation
Determination of Left / Right Simmetry
Neuropeptide Signaling Pathway
Glutamate Receptor Signaling Pathway
Positive Regulation of Synapse Assembly
Nervous System Process
Behavioral Fear Response
G
CTL02A
CTL04E
DAPI
TUJ1
BARHL1
TTR
1st Differentiation
2nd Differentiation
200μm

**Figure S1**

**(A)** Histograms showing Area Under the Curve (AUC) measurements for line CTL02A, CTL04E and CTL08A respectively. Results are displayed in Figure 1 Caption. **(B)** Violin plots showing area measurements for CblOs derived from CTL02A, CTL04E and CTL08A at 3, 7 and 14 days of differentiation. Measurements derive from three different differentiation batches. N = 261, 284 and 265 organoids at 3 days of differentiation for CTL02A, CTL04E and CTL08A respectively; n = 273, 272 and 286 at 7 days and n = 275, 257 and 275 at 14 days. One way ANOVA with Kruskal-Wallis test; * = p<0,0001. **(C)** Representative live imaging pictures of CblOs from CTL08A, acquired with a confocal-spinning disk at 90 days of differentiation with magnification of a cell expressing DLX1/2-GFP (green). Background subtraction was performed with FIJI by using a rolling ball radius of 100. Z-stack of 46. **(D)** Representative pictures of 130 days-old CblOs attached to high density MEA from the control lines used within this study (namely CTL02A, CTL04E and CTL08A), along with representative raster plots showing the inhibitory effect of TTX on spontaneous activity in CblOs derived from the three hiPSC control lines. **(E)** Histograms showing MFR extraburst in CTL02A, CTL04E and CTL08A. Each dot represents the mean value for each active electrode recorded per organoid. Measures coming from two biological replicates, 4-8 differentiation batches CTL02A n = 5, CTL04E n = 9 and CTL08A n = 5. One way ANOVA with Kruskal-Wallis test CTL02A vs. CTL04E p=0,7028; CTL02A vs. CTL08A p= 0,2445; CTL04e vs CTL08A p >0.9999. **(F)** Bubble plot showing the functional characterization of upregulated genes by functional enrichment analysis for the following comparisons: 14 days vs 0 days; 21 days (salmon) vs 14 days (light brown); 28 days vs 21 days (green). The top 8 GO categories from the Biological Process domain of the GO are reported for each comparison (details in methods section). **(G)** Representative confocal-spinning disk images of immunostaining followed by tissue clearing of CblOs from 2 differentiation rounds and 2 cell lines (CTL02A and CTL04E respectively), stained for TTR (red), BARHL1 (green) and TUJ1 (purple). Nuclei were stained with DAPI (blue).
