## Supplementary Figure 2 for "Benchmarking cerebellar organoids to model autism spectrum disorder and human brain evolution"

### Figure S2

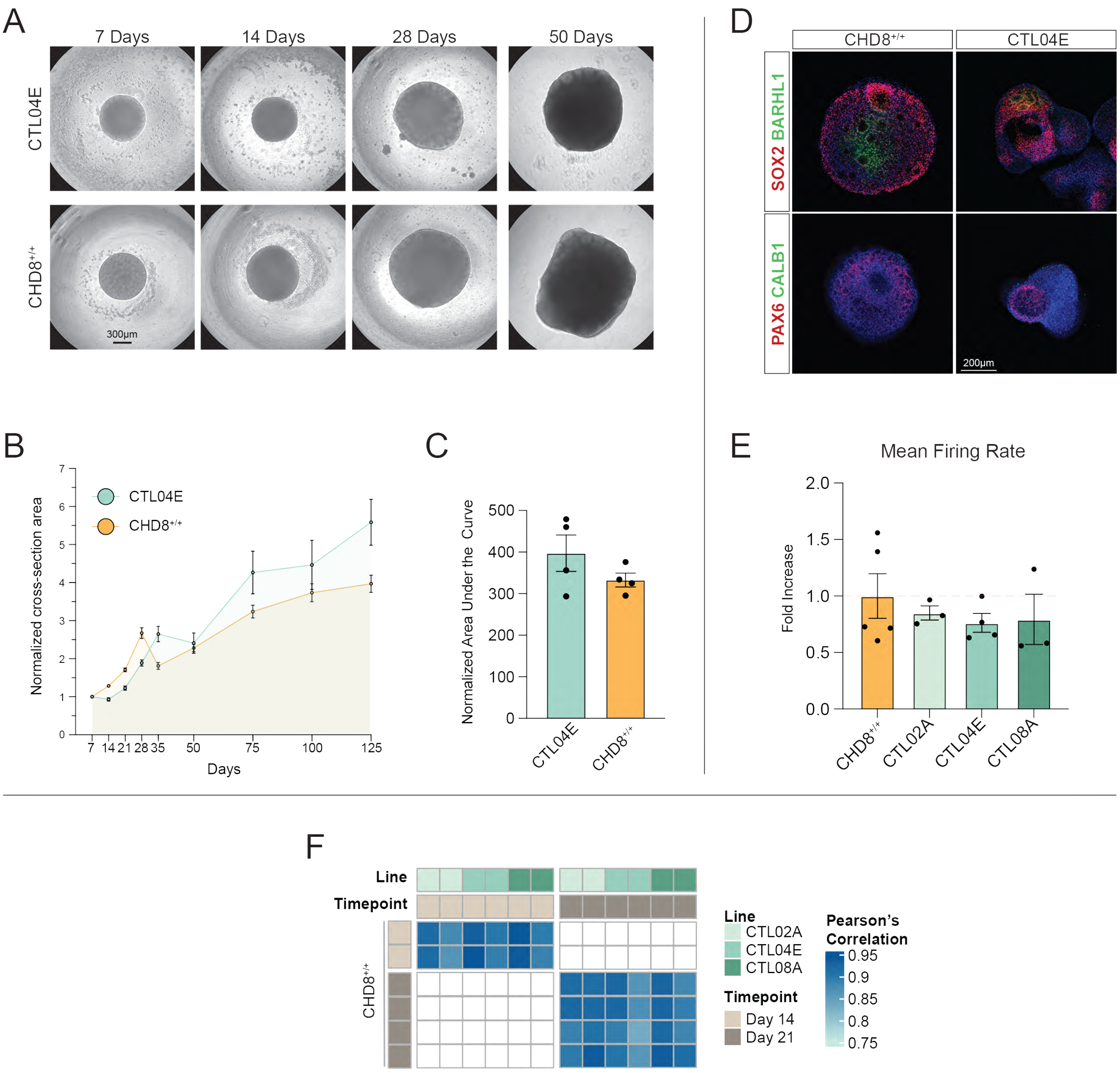
A
7 Days
14 Days
28 Days
50 Days
CTL04E
CHD8+/+
300μm
D
CHD8+/+
CTL04E
SOX2 BARHL1
PAX6 CALB1
200μm
B
CTL04E
CHD8+/+
Normalized cross-section area
Days
C
Normalized Area Under the Curve
CTL04E
CHD8+/+
E
Mean Firing Rate
Fold Increase
CHD8+/+
CTL02A
CTL04E
CTL08A
F
Line
Timepoint
CHD8+/+
Line
CTL02A
CTL04E
CTL08A
Timepoint
Day 14
Day 21
Pearson's Correlation
0.95
0.9
0.85
0.8
0.75

**Figure S2**

**(A)** Representative transmitted light images from CTL04E and CHD8$^{+/+}$ -derived CblOs at 7, 14, 21, 28 and 50 days of differentiation. **(B)** Growth curves comparison from 7 to 125 days of differentiation of CTL04E and CHD8$^{+/+}$ -derived CblOs. **(C)** Histograms representing the AUC calculated on the normalized cross-section area from CHD8$^{+/+}$ and CTL04E. Data are from 4 differentiation round of CblOs from 4-10 organoids per time point. CHD8+/+ =332,6 397,3±16,76 (arbitrary units – a.u.); CTL04E = 397,3± 43,82 (a.u.) p = 0,4857; Mann Whitney test. **(D)** Representative spinning disk confocal images of Cblos from CHD8$^{+/+}$ and CTL04E stained for either SOX2 (green) and BARHL1 (red) or PAX6 (green) and CALB1 (red) at 21 days of differentiation. **(E)** Histograms showing the fold change of the mean firing rate from CHD8$^{+/+}$-derived CblOs compared to organoids from the three hiPSCs control lines (CTL02A, CTL04E and CTL08A respectively). Analysis from two biological replicates, at least 4 differentiation batches: CHD8$^{+/+}$=5 (1,000±0,1973); CTL02A n = 3 (0,8494±0,06230); CTL04E n = 4 (0,7627±0,08342) andCTL08A n = 3 (0,7930±0,2226). ANOVA with Dunn's multiple comparison with respect to CHD8$^{+/+}$: CTL02A p>0.9999, CTL04E p>0.9999, CTL08A p=0,7847. **(F)** Correlation analysis of Cblos derived from hiPSCs (CTL02A, CTL04E and CTL08A respectively) from 0 to 50 days of differentiation *versus* transcriptome from CHD8$^{+/+}$-derived Cblos 14 and 21 days of differentiation. Pearson correlation values (from 0.74 to 0.95), are indicated.
