## Supplementary Figure 3 for "Benchmarking cerebellar organoids to model autism spectrum disorder and human brain evolution"

### Figure S3

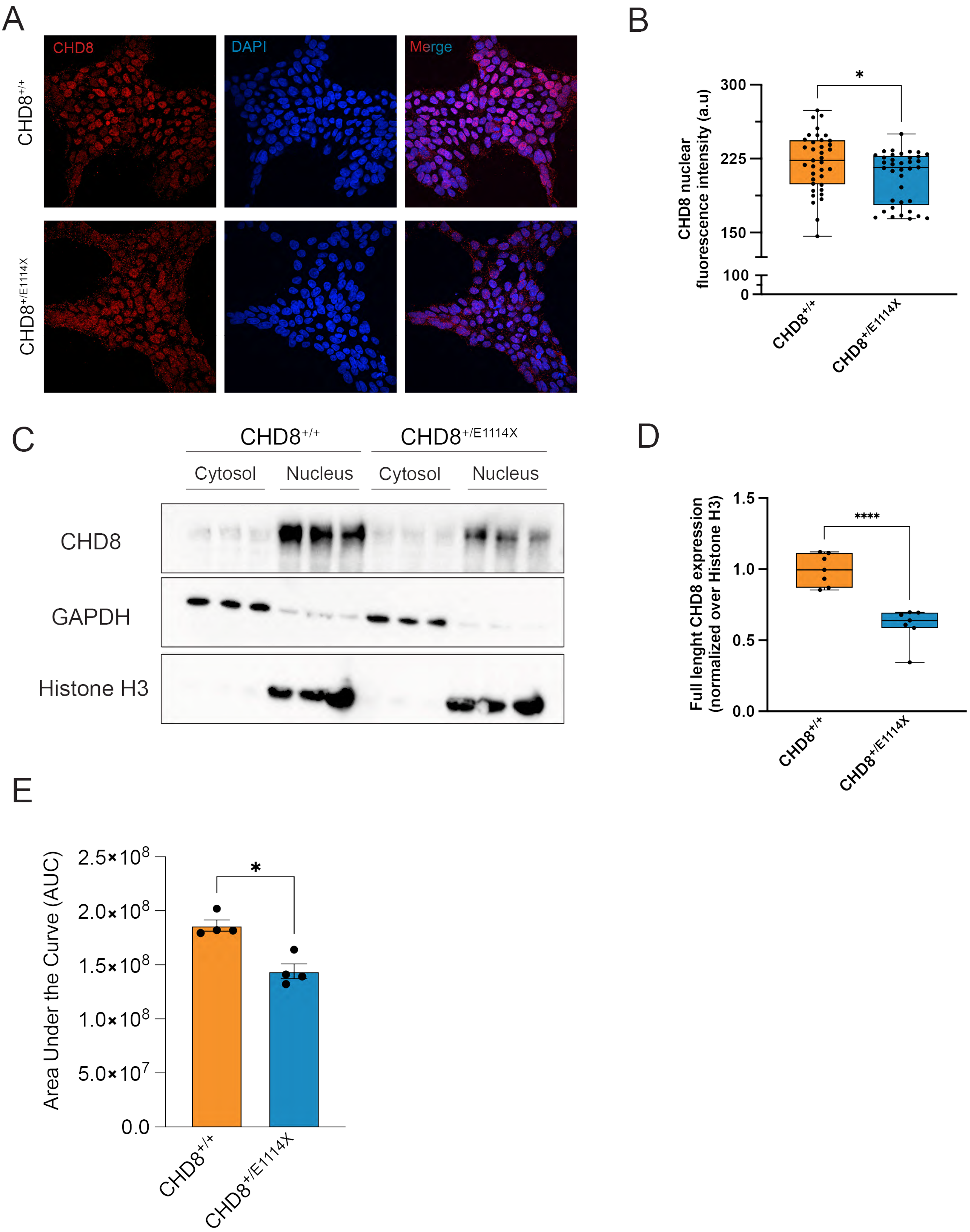

A
CHD8
DAPI
Merge
CHD8+/+
CHD8+/E1114X
B
CHD8 nuclear fluorescence intensity (a.u)
300
225
150
100
0
*
CHD8+/+
CHD8+/E1114X
C
CHD8+/+
CHD8+/E1114X
Cytosol
Nucleus
Cytosol
Nucleus
CHD8
GAPDH
Histone H3
D
Full lenght CHD8 expression (normalized over Histone H3)
1.5
1.0
0.5
0.0
****
CHD8+/+
CHD8+/E1114X
E
Area Under the Curve (AUC)
2.5×10^8
2.0×10^8
1.5×10^8
1.0×10^8
5.0×10^7
0.0
*
CHD8+/+
CHD8+/E1114X

**Figure S3**

**(A)** Representative confocal images from $CHD8^{+/+}$ and $CHD8^{+/E1114X}$ hESCs, stained for CHD8 (red) and DAPI (blue). The fluorescence intensity of CHD8 at nuclear level was performed applying a mask obtained on the DAPI staining and quantified. **(B)** Box plots showing the fluorescence intensity from $CHD8^{+/+}$ = 221,2 ± 4,862 (a.u) and $CHD8^{+/E1114X}$ = 206,4 ± 4,333 (a.u.). Data from 37 fields of view *per* condition from 3 independent experiments. Unpaired t test with Welch's correction; p = 0,0262 (*) **(C)** Representative western blots showing cytosol/nucleus fractionation from $CHD8^{+/+}$ and $CHD8^{+/E1114X}$ hESCs, where GAPDH and Histone H3 were used as cytosolic and nuclear markers respectively. **(D)** Box plot showing the quantification of CHD8 expression in $CHD8^{+/+}$ and $CHD8^{+/E1114X}$ hESCs in the nuclear compartment. $CHD8^{+/+}$ = 0,9942 ± 0,04249; $CHD8^{+/E1114X}$ = 0,6072 ± 0,04662. Data from 7 experimental replicates were analyzed with unpaired t test followed by Welch's correction; p < 0,0001 (****). **(E)** Histograms showing the AUC quantification from $CHD8^{+/+}$ = 186286208±5234036 and $CHD8^{+/E1114X}$ = 144132820±6892421; Mann-Whitney test. (*) p<0,05
