## Supplementary Figure 4 for "Benchmarking cerebellar organoids to model autism spectrum disorder and human brain evolution"

### Figure S4

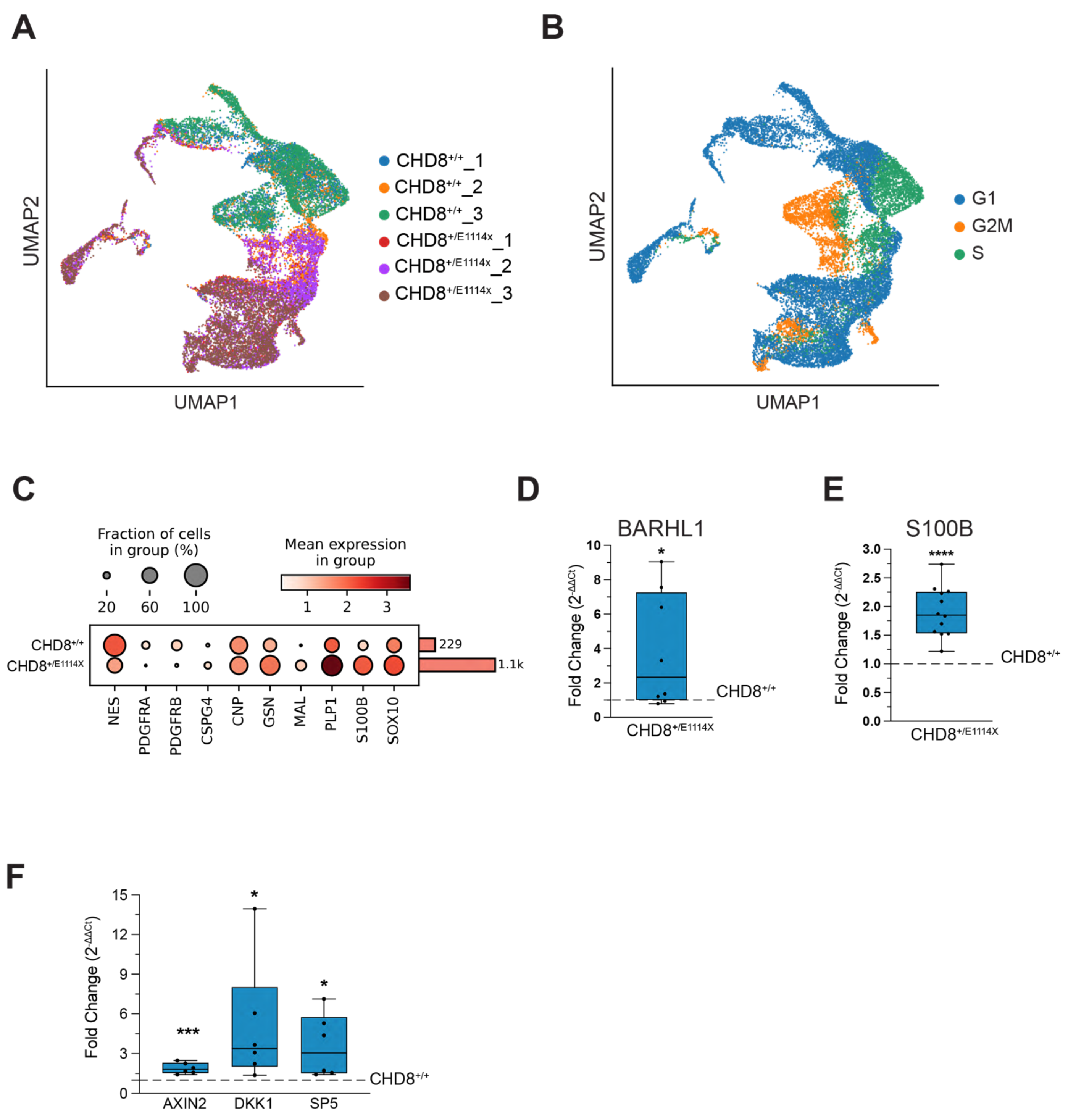


**Figure S4**

**(A-B)** UMAP **c**alculated on the scRNAseq data from CHD8$^{+/E1114X}$ and CHD8$^{+/+}$ CbIOs at day 21 and colored according to experimental replicates (A) or cell-cycle score (B) (G1, G2/M and S phase). **(C)** Bubble plot showing for the same datasets the expression levels of representative marker genes identifying both OPCs (NES, PDGFRA/B, CSPG4) and OL (CNP, GSN, MAL, PLP1, S100B and SOX10) for $CHD8^{+/+}$ and $CHD8^{+/E1114X}$. **(D)** Box plot showing qPCR measuring BARHL1 expression in $CHD8^{+/E1114X}$-derived organoids at 21 days of differentiation. Fold change measured upon normalization of $2^{-\Delta\Delta Ct}$ over control $CHD8^{+/+}$. Paired *t*-test calculated on ΔCt from the two conditions. p=0,0126 (*) n=8 from 4 independent rounds of differentiation. **(E)** Box plot showing qPCR measuring S100B expression in $CHD8^{+/E1114X}$-derived organoids at 21 days of differentiation. Fold change measured upon normalization of $2^{-\Delta\Delta Ct}$ over control $CHD8^{+/+}$. Paired *t*-test calculated on ΔCt from the two conditions, p=0,0293 (*). n=12 from 4 independent rounds of differentiation. **(F)** Box plot showing gene expression of markers involved in b-catenin repression from CblOs at 21 days of differentiation. Values are expressed as fold change of $CHD8^{+/E1114X}$ normalized on $CHD8^{+/+}$. Paired *t* test on ΔΔCt was used. AXIN2 p=0,0009 (***), DKK1 p=0,0102 (*), SP5 p=0,0139 (*).
