## Supplementary Figure 5 for "Benchmarking cerebellar organoids to model autism spectrum disorder and human brain evolution"

### Figure S5

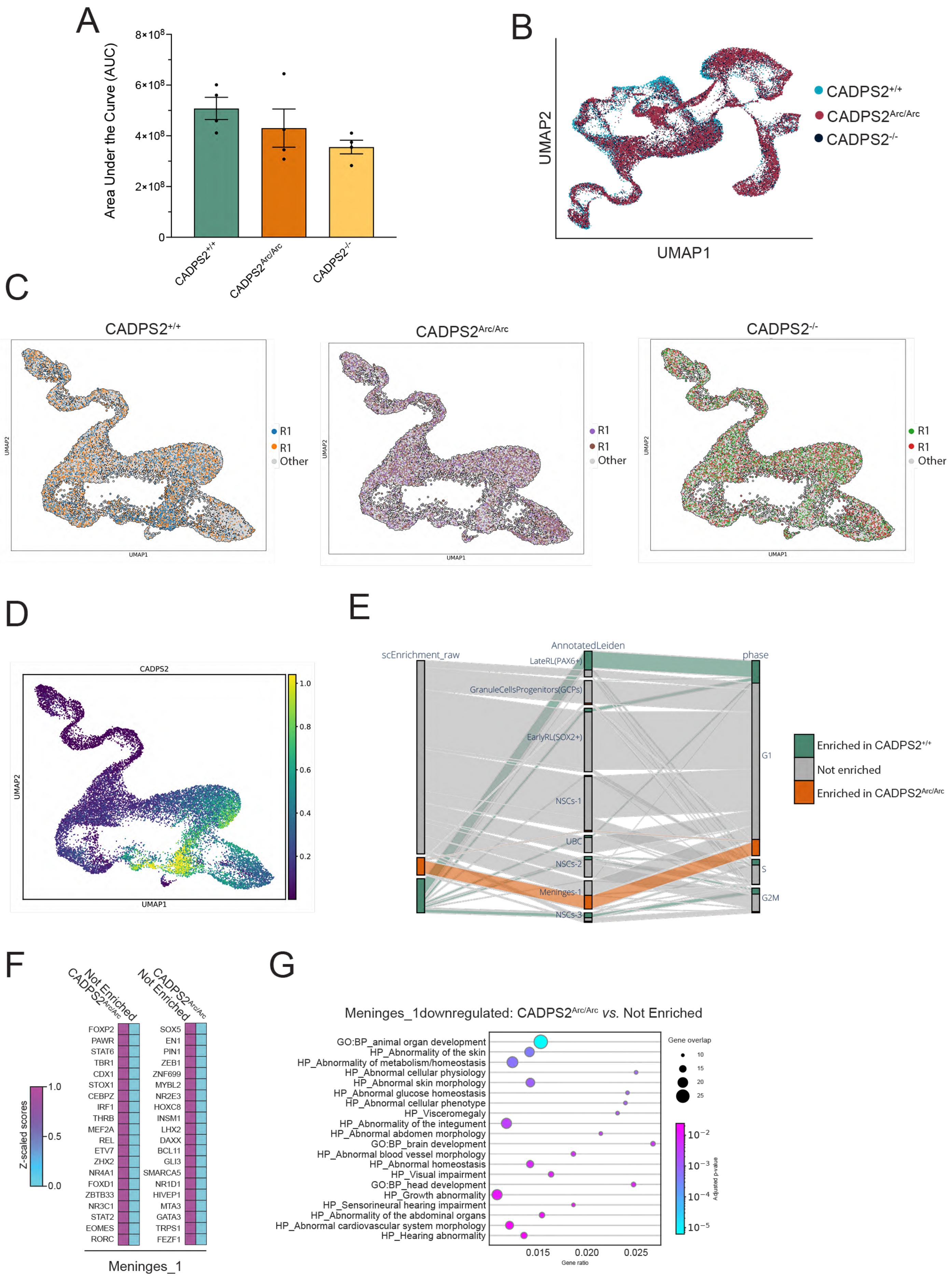

A
Area Under the Curve (AUC)
8×10^8
6×10^8
4×10^8
2×10^8
0
CADPS2+/+
CADPS2Arc/Arc
CADPS2-/-
B
UMAP2
UMAP1
CADPS2+/+
CADPS2Arc/Arc
CADPS2-/-
C
CADPS2+/+
CADPS2Arc/Arc
CADPS2-/-
R1
R1
Other
UMAP1
UMAP2
D
CADPS2
UMAP1
UMAP2
1.0
0.8
0.6
0.4
0.2
E
scEnrichment_raw
AnnotatedLeiden
phase
LateRL(PAX6+)
GranuleCellsProgenitors(GCPs)
EarlyRL(SOX2+)
NSCs-1
UBC
NSCs-2
Meninges-1
NSCs-3
G1
S
G2M
Enriched in CADPS2+/+
Not enriched
Enriched in CADPS2Arc/Arc
F
CADPS2Arc/Arc
Not Enriched
Not Enriched
CADPS2Arc/Arc
Z-scaled scores
1.0
0.5
0.0
FOXP2
PAWR
STAT6
TBR1
CDX1
STOX1
CEBPZ
IRF1
THRB
MEF2A
REL
ETV7
ZHX2
NR4A1
FOXD1
ZBTB33
NR3C1
STAT2
EOMES
RORC
SOX5
EN1
PIN1
ZEB1
ZNF699
MYBL2
NR2E3
HOXC8
INSM1
LHX2
DAXX
BCL11
GLI3
SMARCA5
NR1D1
HIVEP1
MTA3
GATA3
TRPS1
FEZF1
Meninges_1
G
Meninges_1downregulated: CADPS2Arc/Arc vs. Not Enriched
GO:BP_animal organ development
HP_Abnormality of the skin
HP_Abnormality of metabolism/homeostasis
HP_Abnormal cellular physiology
HP_Abnormal skin morphology
HP_Abnormal glucose homeostasis
HP_Abnormal cellular phenotype
HP_Visceromegaly
HP_Abnormality of the integument
HP_Abnormal abdomen morphology
GO:BP_brain development
HP_Abnormal blood vessel morphology
HP_Abnormal homeostasis
HP_Visual impairment
GO:BP_head development
HP_Growth abnormality
HP_Sensorineural hearing impairment
HP_Abnormality of the abdominal organs
HP_Abnormal cardiovascular system morphology
HP_Hearing abnormality
0.015
0.020
0.025
Gene ratio
Gene overlap
10
15
20
25
Adjusted p-value
10^-2
10^-3
10^-4
10^-5

**Figure S5**

**(A)** Histograms showing the AUC from CADPS2$^{+/+}$, CADPS2$^{Arc/Arc}$ and CADPS2$^{-/-}$-derived CblOs at 125 days of differentiation. Results from 4 rounds of differentiation. One way anova with Kruskal-Wallis test: CADPS2$^{+/+}$ 507940348±43937614; CADPS2$^{Arc/Arc}$ 430438885±75403503; CADPS2$^{-/-}$ 355850988± 26901247. CADPS2$^{+/+}$ vs. CADPS2$^{Arc/Arc}$ p= 0,7179 (ns); CADPS2$^{+/+}$ vs. CADPS2$^{-/-}$ p= 0,1184 (ns); CADPS2$^{Arc/Arc}$ vs. CADPS2$^{-/-}$ p= >0.9999 (ns). **(B)** UMAP embeddings colored by genotype in scRNAseq experiments from CblOs at 25 days of differentiation. Genotypes displayed CADPS2$^{+/+}$ (light blue), CADPS2$^{Arc/Arc}$ (red purple) and CADPS2$^{-/-}$ (black). **(C)** UMAP embeddings colored by experimental replicates (R1 and R2) of each genotype. Cells belonging to other genotypes are colored in grey. **(D)** UMAP showing the expression levels and distribution of CADPS2 in CADPS2-expressing cells and their closest derivatives. Expression levels are displayed as MAGIC imputed counts[1] with values trimmed between 1$^{st}$ and 99$^{th}$ percentile. **(E)** Sankey diagram showing the mapping of cells in CADPS2$^{+/+}$ or CADPS2$^{Arc/Arc}$ enriched neighborhood to annotated clusters and cell cycle phases. The plot shows the mapping for both differentially abundant and not differentially abundant cell types (Star methods). **(F)** TF activity analysis performed on the differential abundance clustering between CADPS2$^{Arc/Arc}$ enriched *vs.* Not enriched domains in meninges. **(G)** Functional enrichment of DEGs deriving from CADPS2$^{Arc/Arc}$ enriched *vs.* Not enriched domains in cells from the meninges cluster.

1. van Dijk, D., Sharma, R., Nainys, J., Yim, K., Kathail, P., Carr, A.J., Burdziak, C., Moon, K.R., Chaffer, C.L., Pattabiraman, D., et al. (2018). Recovering Gene Interactions from Single-Cell Data Using Data Diffusion. Cell *174*, 716-729.e27. https://doi.org/10.1016/J.CELL.2018.05.061.
