## Supplementary Figure 5 for "Benchmarking cerebellar organoids to model autism spectrum disorder and human brain evolution"

### Figure S6

A

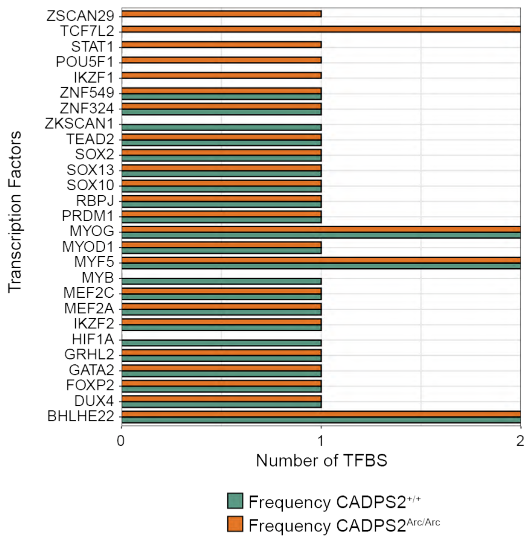


B

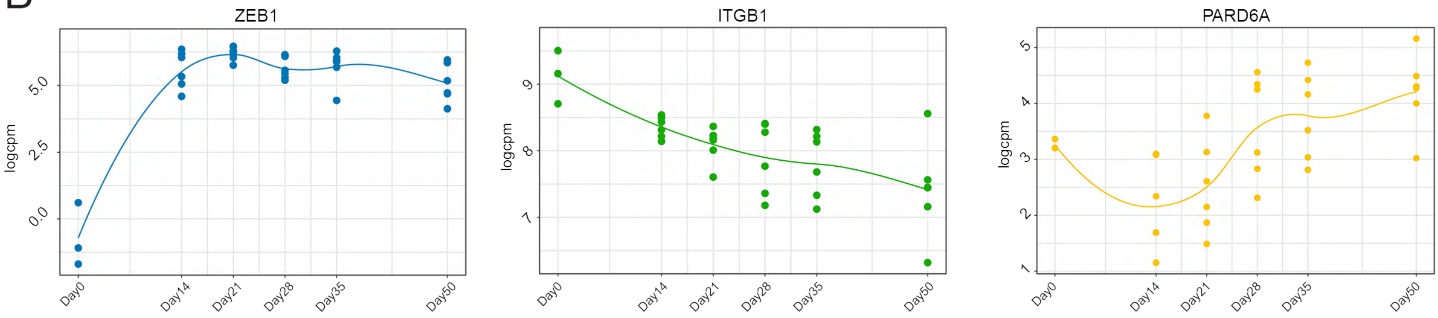


**Figure S6**

**(A)** Frequency of all transcription factors binding site predicted in the 37bp window centered on the SNV. **(B)** Trend plots showing the expression levels (in logcpm) of ZEB1, ITGB1 and PARD6A in our longitudinal bulk RNASeq dataset from CblOs at 0, 14, 21, 28, 35 and 50 days of differentiation. Each line represents interpolated values from each dot, from the three control lines CTL02A, CTL04E and CTL08A (two replicates per cell line).
