## Supplementary material for "Benchmarking cerebellar organoids to model autism spectrum disorder and human brain evolution": Table S1

- qPCR Oligonucleotides

| Gene | Forward (F) / Reverse (R) | Sequence 5'-3' |
|---|---|---|
| AXIN2 | F | GAGTGGACTTGTGCCGACTTCA |
| | R | GGTGGCTGGTGCAAAGACATAG |
| BARHL1 | F | GAGCGGCAGAAGTACCTGAG |
| | R | GTAGAAATAAGGCGACGGGAAC |
| DKK1 | F | GATCATAGCACCTTGGATGGG |
| | R | GGCACAGTCTGATGACCGG |
| S100B | F | TGGCCCTCATCGACGTTTTC |
| | R | ATGTTCAAAGAACTCGTGGCA |
| SP5 | F | GTCCTCATCGTCGTGGTGATTG |
| | R | AGAAGGTGGCAGTGGTAACCAG |
| TBP | F | GCCACGCCAGCTTCGGAGAG |
| | R | CCGCAGCAAACCGCTTGGGA |

- sgRNA: CADPS2$^{-/-}$

| Gene | Forward (F) / Reverse (R) | Sequence 5'-3' |
|---|---|---|
| sgRNA1 | F | CACCAGCGAAGAGGAGTCGGACGA |
| | R | AAACTCGTCCGACTCCTCTTCGCT |

- sgRNA: CADPS2$^{Arc/Arc}$

| Gene | Forward (F) / Reverse (R) | Sequence 5'-3' |
|---|---|---|
| sgRNA_SNP1 | F | CACCAATAAGGTCTGAAGCATACA |
| | R | AAACTGTATGCTTCAGACCTTATT |

- ssODN

| Gene | Genomic locus | Sequence 5'-3' |
|---|---|---|
| SNP-1 | (chr7: 122'715'624) Ch38/hg38 | GACTCTCTTTTCTGTACACTGTCCAAACAGAGGTCATTCTGCTTTG AATAAGGTCTGAAGCATACAGGGTCTGCCTGTAAAACGAAAGCT GCTTTCCTATGGACAAAGATAGCTTTAGTCATCTTCTATGGTCAA AATTCTCACAGACACAAGGCTTAAAACCTCTCAACACAACGCATT GCAAAGAAAAAAAAAAAGCC |

- Primers Sanger sequencing

| Gene | Forward (F) / Reverse (R) | Sequence 5'-3' |
|---|---|---|
| CADPS2$^{-/-}$ | F | GGCTCTTCAACATTCAATTC |
| | R | AAATTAAAGTTGATGGGAGT |
| CADPS2$^{Arc/Arc}$ | F | ACCTTACTAGGGGCAGACCA |
| | R | GGCAAAGTGGTGACTTGAGC |
